# Excessive self-grooming, gene dysregulation and imbalance between the striosome and matrix compartments in the striatum of *Shank3* mutant mice

**DOI:** 10.1101/2022.01.19.476922

**Authors:** Ferhat Allain-Thibeault, Verpy Elisabeth, Biton Anne, Forget Benoît, Fabrice de Chaumont, Mueller Florian, Le Sourd Anne-Marie, Coqueran Sabrina, Schmitt Julien, Rochefort Christelle, Rondi-Reig Laure, Leboucher Aziliz, Boland Anne, Fin Bertrand, Deleuze Jean François, Tobias M. Boeckers, Ey Elodie, Bourgeron Thomas

**Affiliations:** Génétique Humaine et Fonctions Cognitives, Institut Pasteur, UMR3571 CNRS, Université de Paris Cité, Paris, France; Department of Neuroscience, Columbia University Medical Center, New York, NY, 10027; Zuckerman Mind Brain Behavior Institute, Columbia University, New York, NY, 10027; Centre de Bioinformatique, Biostatistique et Biologie Intégrative (C3BI, USR 3756 Institut Pasteur and CNRS), Paris, France; Unité Imagerie et Modélisation (UMR 3691 Institut Pasteur and CNRS), Paris, France; Sorbonne Université, CNRS, INSERM, Institut de Biologie Paris Seine (IBPS), Neurosciences Paris Seine (NPS), Cerebellum Navigation and Memory Team (CeZaMe), F-75005 Paris, France; Université Paris-Saclay, CEA, Centre National de Recherche en Génomique Humaine (CNRGH), 91057, Evry, France; Centre d’étude du polymorphisme humain, Paris, France; Institute of Anatomy and Cell Biology, Ulm University, 89081 Ulm, Germany; Deutsches Zentrum fur Neurodegenerative Erkrankungen (DZNE), 89081 Ulm, Germany; Université de Strasbourg, CNRS, INSERM, Institut de Génétique et de Biologie Moléculaire et Cellulaire - UMR 7104, Illkirch-Graffenstaden, France

## Abstract

Autism is characterised by atypical social communication and stereotyped behaviours. Mutations in the gene encoding the synaptic scaffolding protein SHANK3 are detected in 1-2% of patients with autism and intellectual disability (ID), but the mechanisms underpinning the symptoms remain largely unknown. Here, we characterised the behaviour of *Shank3*^Δ11/Δ11^ mice from three to twelve months of age. We observed decreased locomotor activity, increased stereotyped self-grooming and modification of socio-sexual interaction compared to wild-type littermates. We then used RNAseq on four brain regions of the same animals to identify differentially expressed genes (DEG). DEGs were identified mainly in the striatum and were associated with synaptic transmission (e.g. *Grm2, Dlgap1*), G-protein-signalling pathways (e.g. *Gnal, Prkcg1, and Camk2g*), as well as excitation/inhibition balance (e.g. *Gad2*). Downregulated and upregulated genes were enriched in the gene clusters of medium-sized spiny neurons expressing the dopamine 1 (D1-MSN) and the dopamine 2 receptor (D2-MSN), respectively. Several DEGs (*Cnr1, Gnal1, Gad2, and Drd4*) were reported as striosome markers. By studying the distribution of the glutamate decarboxylase GAD65, encoded by *Gad2*, we showed that the striosome compartment of *Shank3*^Δ11/Δ11^ mice was enlarged and displayed much higher expression of GAD65 compared to wild-type mice. Altogether, these results indicate altered gene expression in the striatum of SHANK3-deficient mice and strongly suggest, for the first time, that the impairment in behaviour of these mice are related to an imbalance striosomes/matrix.

## Introduction

Autism is a neurodevelopmental condition characterised by atypical social communication and interactions associated with stereotyped behaviours and restricted interests. More than 200 genes have been robustly associated with autism pointing at biological pathways such as chromatin remodelling, protein translation and synaptic function ^1^. Among these genes, *SHANK3* (SH3 and multiple ankyrin repeat domain 3) codes for a scaffolding protein located at the postsynaptic density (PSD) of glutamatergic synapses where it interacts with other scaffolding proteins, cytoskeletal proteins, glutamate receptors and signalling molecules ^2^.

Heterozygous *de novo* mutations deleting the *SHANK3* gene on chromosome 22q13.3 or affecting its coding region are associated with Phelan-McDermid syndrome ^3^ and observed in 1-2% of patients with both autism and intellectual disability (ID), making this gene a major cause of neurodevelopmental disorder (NDD) ^4^. Remarkably, a relatively high proportion of patients (>60%) carrying *SHANK3* mutations display regression (i.e., substantial loss of language and social skills) during adolescence and adulthood ^5,6^.

Through the use of multiple intragenic promoters and alternative splicing, several SHANK3 protein isoforms are differentially expressed according to developmental stage or brain region ^7^. Numerous *Shank3* mutant mice have been generated [for reviews see ^2,8^]. Most of them bear deletion of specific exons still allowing the expression of some isoforms ^7,9^. The behavioural deficits reported for *Shank3* mutant mice depend on the strains and experimental conditions. However, the presence of stereotyped-like behaviours, such as excessive self-grooming, was reported in all *Shank3* mutant mice and is reminiscent of the stereotyped behaviours observed in patients with autism ^2,10,11^. Stereotyped behaviours are frequently associated with anomalies of the striatum ^12^, a region that coordinates multiple aspects of cognition, including action planning, decision-making, motivation, reinforcement, and reward perception as well as motor control. Several studies identified anomalies in the striatum of *Shank3* mutant mice at the anatomical (e.g., larger volume), circuit (e.g., disruption of indirect pathway of basal ganglia) and cellular (e.g., decreased striatal expression of GluN2B, a NMDA receptor subunit) levels ^13–18^. The biological pathways underpinning the observed abnormalities remain however largely unknown.

In the present work, we first conducted a thorough longitudinal behavioural characterization of *Shank3*^Δ11/Δ11^ mice, carrying a deletion of exon 11 of *Shank3* ^19^, at three, eight and twelve months of age, focusing on social interactions, locomotion and stereotyped behaviours. We then compared gene expression in four brain structures - cortex, hippocampus, cerebellum and striatum - in *Shank3*^*+/+*^ and *Shank3*^Δ11/Δ11^ male littermates characterised in the behavioural study. Compared to the other brain regions, the striatum appeared to be highly sensitive to the loss of SHANK3. We also identified a correlation between the level of self-grooming and an increased striatal expression of several genes, including *Gad2* encoding GAD65, which catalyses the transformation of glutamate into GABA at the synapse. Finally, using immunofluorescence on striatum sections at 12 months, we found that, compared to wild-type littermates, the *Shank3*^Δ11/Δ11^ mice displayed an enlargement of the compartment formed by striosomes/patches which also markedly over-expressed GAD65. These alterations of the striosomal compartment were also found in the *Shank3*^Δ4-22/Δ4-22^ mouse with a complete lack of all SHANK3 isoforms (deletion from exon 4 to 22). Altogether our results point toward a correlation between excessive self-grooming and signalling imbalance between the striosome and matrix compartments of the striatum in SHANK3-deficient mice.

## Results

At three months of age, *Shank3*^Δ11/Δ11^ mice, from Cohort 1 and 2, displayed typical body weight, anxiety-like behaviour levels and working memory, compared to *Shank3*^+/+^ littermates (**Fig. S1**). Considering the frequent regression of patients carrying *SHANK3* mutations, we tested whether a phenotypic deterioration occurred in *Shank3*^Δ11/Δ11^ mice. For that purpose, we tested the main impairments observed in the *Shank3*^Δ11^ mouse model over time, at three, eight and twelve months of age: exploratory activity, social communication and stereotyped behaviours in mice.

### Hypoactivity and atypical motor abilities in *Shank3*^Δ11/Δ11^ mice

Impairment in locomotor activities is often observed in *Shank3* mutant mice (Ferhat et al., 2017). In the open field at three months of age, *Shank3*^Δ11/Δ11^ males travelled significantly shorter distances in comparison with *Shank3*^+/+^ littermates (Cohort 1: W=130, p=0.007; **Fig. 1A**, left panel). This observation was not significant for *Shank3*^Δ11/Δ11^ females of Cohort 1 but was confirmed for *Shank3*^Δ11/Δ11^ males of Cohort 2 (**Fig. S2A**) and Cohort 3 (**Fig. S2B**) at three months of age. This reduction of distance travelled in *Shank3*^Δ11/Δ11^ males was not present at older age; however, *Shank3*^+/+^ and *Shank3*^+/Δ11^ mice reduced their activity with increasing age (**Fig. 1A**). A similar trend was present but did not reach significance levels in females (Cohort 1: W=92, p=0.07; **Fig. 1A**). Remarkably, in stressful swimming conditions, *Shank3*^Δ11/Δ11^ mice, from Cohort 3, swam at a significantly higher speed compared to wild-type mice in the Morris water maze with visible platform (Cohort 3: p<0.01 over all training) and in the star maze (p<0.01), without any significant difference to find the direct path in the star maze (**Fig. S2E-G**). *Shank3*^Δ11/Δ11^ males also spent less time digging in the bedding in comparison with *Shank3*^+/+^ mice at three months (Cohort 1: W=123, p=0.027; **Fig. 1B;** Cohort 2: W=173, p<0.001; **Fig. S2C**). Genotype-related differences in digging behaviour persisted with increasing age (8 months: W= 140, p<0.01; 12 months; W=107; p=0.01; **Fig. 1B**).

**Figure 1:**
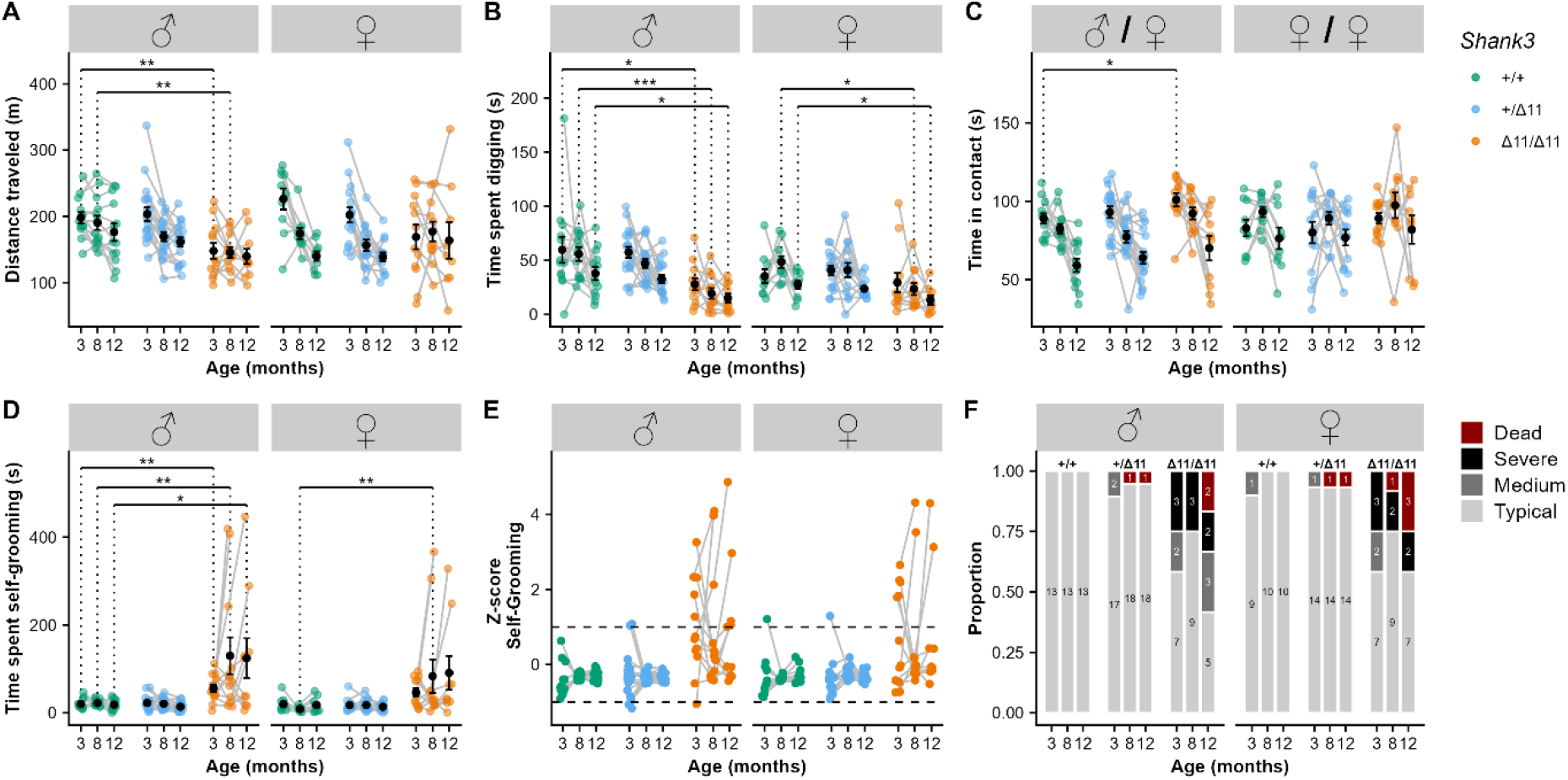
Hypoactivity, reduced exploration and increased stereotyped behaviours in *Shank3*^Δ11/Δ11^ mice at three, eight and twelve months of age. (A) Total distance travelled during 30-min free exploration of an open field in wild-type (green), heterozygous (blue) and *Shank3*^Δ11/Δ11^ (orange) for males (left panel) and females (right panel). (B) Total time spent digging in fresh bedding during 10 min observation in a new test cage, after 10 min habituation. (C) Total time spent in contact during male/female and female/female social interaction. (D) Total time spent self-grooming during 10 min observation in a new test cage, after 10 min habituation. (E) Z-score for the time spent self-grooming. (F) Proportion and number of individuals displaying different levels of severity in self-grooming. “Dead” corresponds to animals that had to be euthanized due to severe self-inflicted injuries. A-D: Black points represent mean with standard error of the mean (s.e.m). Mann–Whitney U test with Bonferroni correction for multiple testing (2 tests): *: corrected p-value <0.05; **: corrected p-value <0.01.

### Atypical socio-sexual behaviour in *Shank3*^Δ11/Δ11^ male mice at early stage

Impairments in social interactions and communication, one of the core symptoms of autism, were not always observed in the different *Shank3* mouse models ^8^. We analysed the preference for a conspecific during three-chambered test and the time of interaction and types of contact during free dyadic interaction (**Fig. S3**). During the three-chambered test, mice of all genotypes and both sexes displayed a typical preference for a same-sex conspecific (**Fig. S4**). Furthermore, no significant difference was observed during free dyadic social interaction, at different ages for Cohort 1, in the latency for the first contact (not shown), in the emission of ultrasonic vocalisations (USVs) **(Fig. S4B)**, and in the total time spent in contact and in the different social behaviours with same sex conspecific for both sexes (**Fig. S3, S4 C-F**).

In contrast, we observed that *Shank3*^Δ11/Δ11^ males spent significantly more time in contact with an oestrous female compared to *Shank3*^+/+^ males at three months of age (W=32, p=0.034; **Fig. 1C**), especially in oro-oral contact (W=22, p=0.005; **Fig. S3, S4G**). We also observed a reduction of time keeping the female in the visual field for *Shank3*^Δ11/Δ11^ males compared to *Shank3*^+/+^ males (W=133, p=0.006; **Fig. S4G**). These altered behaviours were not accompanied with atypical vocal behaviour (**Fig. S4B**). At later age, the increase of overall social interaction disappeared during interaction between *Shank3*^Δ11/Δ11^ males and estrous females. However, at eight months of age, the duration of oro-oral contact in *Shank3*^Δ11/Δ11^ males with an oestrous female was still significantly increased (W=29, p=0.025), but these differences were no longer significant at twelve months of age (**Fig. S4H-I**). Nevertheless, at this age, we detected an increase of the “approach then escape’’ behavioural sequence displayed by *Shank3*^Δ11/Δ11^ males toward a C57BL/6J female in comparison with *Shank3*^+/+^ males (W=15.5, p=0.005, **Fig. S4I**).

### Shank3 mutant mice display excessive self-grooming behaviour with age

Self-grooming is a spontaneous and natural behaviour displayed by mice for hygienic purposes or in reaction to stressful conditions ^12^. Here, we investigated whether this behaviour was exacerbated in mutant mice compared to WT littermates in an unfamiliar test environment. At 3 months of age, *Shank3*^Δ11/Δ11^ male mice already displayed a significant increase in the time spent self-grooming in comparison with *Shank3*^+/+^ males for Cohort 1 (W=21, p=0.002) and 2 (W=43, p=0.022). Similar trends did not reach significance in female mice (**Fig. 1D-F, Fig. S2D**). Furthermore, a subsample of *Shank3*^Δ11/Δ11^ males (5 individuals over 12) and females (6 individuals over 12) presented hair removal that evolved into self-injuries.

At eight and twelve months of age, Cohort 1 *Shank3*^Δ11/Δ11^ males still displayed increased self-grooming in comparison with *Shank3*^+/+^ mice (eight months: W=27, p=0.009; twelve months: W=37, p=0.041; **Fig. 1D**). The same was true for *Shank3*^Δ11/Δ11^ females who also groomed themselves significantly more than *Shank3*^+/+^ females (eight months: W=11, p=0.002; twelve months: W=20, p=0.014; **Fig. 1D**). With age, an increasing number of *Shank3*^Δ11/Δ11^ mice were outliers, performing self-grooming during a period between one and two standard deviation(s) (medium phenotype) or exceeding two standard deviations (severe phenotype) or had to be euthanized because of severe self-inflicted injuries. Hereafter, we denominate mice with a period of self-grooming over one standard deviation as excessive self-groomers. Nevertheless, for 4 out of 12 *Shank3*^Δ11/Δ11^ males and 3 out of 12 *Shank3*^Δ11/Δ11^ females, we did not observe excessive self-grooming at any of the three time points (**Fig. 1D, E**).

### Differential expression of genes related to synaptic function, signalling and cytoskeleton dynamics in the striatum of *Shank3*^Δ11/Δ11^ mice

To investigate the impact of the loss of the major SHANK3 isoforms on the transcriptome, we performed RNAseq on four brain structures (whole cortex, striatum, hippocampus, and cerebellum) of seven *Shank3*^+/+^ and eight *Shank3*^Δ11/Δ11^ male mice after their behavioural characterization at twelve months of age. We observed a reduction of the transcript levels of the major isoforms of *Shank3* in *Shank3*^Δ11/Δ11^ (**Fig. S5**). We also observed an enrichment of DEGs in the vicinity of *Shank3* locus on chromosome 15 (before: 10.4 Mb; after: 7.5 Mb) (**Table S2**). This enrichment is most likely due to the residual genomic region from the 129S1/SvImJ ES cells used to generate the *Shank3*^Δ11/Δ11^ mice (**Fig. S6**) ^19^. We therefore excluded these genes from the analyses (more details in **Supplementary methods**). As expected, *Shank3* expression levels in all four brain structures were significantly reduced in *Shank3*^Δ11/Δ11^ mice in comparison with *Shank3*^+/+^ mice (FDR < 0.001; **Fig. S7**). Notably, the *Shank3* sequence reads still present in the *Shank3*^Δ11/Δ11^ mice were aligned to exons downstream of exon 11 (**Table S2**). No significant changes in the expression of *Shank1* and *Shank2* were found (**Fig. S7, Table S2**).

We examined the DEGs (FDR < 0.05) within each of the four brain structures. The largest number of DEGs was detected in the striatum with 140 DEGs. The hippocampus displayed 26 DEGs, the cortex 13 and the cerebellum 16 (**Fig. 2A**). After removing the genes and pseudogenes located around the *Shank3* locus, *Shank3* was the only gene differentially expressed in the four brain structures (**Fig. 2A**). The DEGs in the striatum displayed an increased dispersion compared to the other structures, indicating an increased variability of response between animals (**Fig. S8**). Altogether, these results suggest that, among the four structures studied, the transcriptome of the striatum is by far the most impacted by the loss of the major isoforms of SHANK3. We performed quantitative RT-PCR (q-RT-PCR) to confirm up- or downregulation of 30 DEGs (Spearman correlation R=0.809 P<0.0001; **Supplementary Fig. S9**).

**Figure 2:**
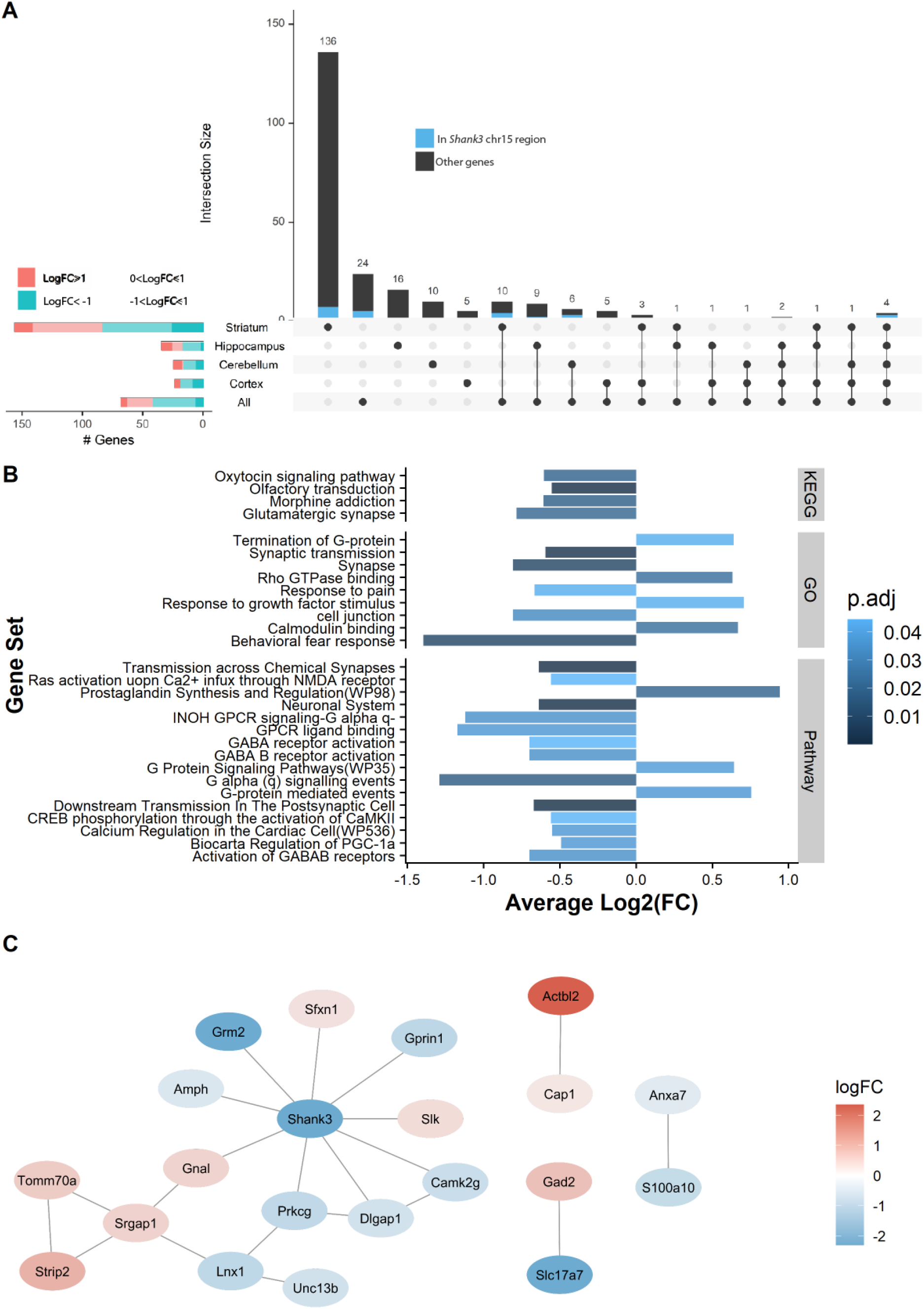
The striatum displays the largest gene expression differences between *Shank3*^+/+^ and *Shank3*^Δ11/Δ11^ mice and the largest number of impacted pathways. (A) Upset plot displaying the intersection between brain structures for the genes differentially expressed between *Shank3*^Δ11/Δ11^ and *Shank3*^+/+^ mice (adjusted p-value < 0.05; determined by limma-voom using observation quality weights). Genes located around the Shank3 region on chromosome 15 are shown in blue. The horizontal bar plots on the left show the number of differentially expressed genes in the different brain structures according to the value of their log-fold changes (logFC). (B) GO (Gene Ontology), GeneSetDB Pathway (gsdbpath), and KEGG (Kyoto Encyclopaedia of Genes and Genomes) gene sets found to be over-represented (top 20 gene sets with adjusted p-value < 0.05) in the genes detected as differentially expressed in the striatum. (C) Protein-protein interaction (PPI) network for the differentially expressed genes (DEGs). Nodes, coloured ovals, represent DEGs; edges, black lines, represent direct protein-protein interactions (from BioGrid) between DEGs products. The colour of the node represents the Log2FC of DEGs: red means up-regulation and blue means down-regulation in Shank3^Δ11/Δ11^ mice compared to wild-type mice.

We investigated the functions of the proteins encoded by the DEGs in the striatum using over-representation analysis (ORA) and Protein-Protein Interaction (PPI) analysis. The ORA (**Table S3**) revealed enrichment in DEGs associated with, among others, two pathways of interest: synaptic transmission (negative fold-change in *Shank3*^Δ11/Δ11^ mice in comparison with *Shank3*^+/+^ mice), and G-protein activity (positive and negative fold-change in *Shank3*^Δ11/Δ11^ mice in comparison with *Shank3*^+/+^ mice according to the type of G-protein pathway) (**Fig. 2B** and **Table S3**). PPI analysis of the striatal DEGs highlighted one network of 15 proteins (p-value from randomised network: <0.01; **Fig. 2C**), some of them directly interacting with SHANK3 in the PSD, such as the metabotropic glutamate receptor 2 (encoded by *Grm2*, highly decreased) and Disks large-associated protein 1, a scaffolding protein (encoded by *Dlgap1*, also decreased) interacting with AMPA and NMDA receptors. In addition to these synaptic proteins, the network contains proteins involved in signal transduction (e.g. *Prkcg1, Camk2g*) and cytoskeleton dynamics (e.g. *Slk*). Other small networks identified by the PPI analysis contain proteins involved in glutamate transportation (*Slc17a7*) and decarboxylation (*Gad2*), and cytoskeleton dynamics (*Actbl2*).

### Cellular expression pattern of the DEGs expressed in the striatum of *Shank3*^Δ11/Δ11^ mice

The inhibitory medium-sized spiny neurons (MSN) constitute the major type of striatal neuronal population. They receive glutamatergic inputs and are the target of dopamine innervation from cortex, thalamus, amygdala and substantia nigra pars compacta (SNpc) ^20^. Most of these MSN express either dopamine 1 receptor DRD1 (D1-MSN, belonging to the direct pathway), dopamine 2 receptor DRD2 (D2-MSN, belonging to the indirect pathway), or both receptors for a small fraction of MSN. In order to identify the striatal cell types which are the most impacted by SHANK3 deficiency, we compared the expression pattern of the DEGs with marker genes of striatal cell types reported in two single-cell RNA sequencing studies ^21,22^. We observed that DEGs under-expressed in *Shank3*^Δ11/Δ11^ compared to *Shank3*^+/+^ mice were enriched in the D1-MSN clusters while overexpressed genes were enriched in the D2-MSN clusters (**Fig. 3A, Table S2 & S4**). This could reflect a modified proportion of D1-MSN (decrease) and D2-MSN (increase) in the striatum of the *Shank3*^Δ11/Δ11^ mice. However, many genes commonly considered as specific to D1-MSN (*Drd1, Isl1, Sfxn1* and *Tac1*) or D2-MSN (*Drd2, Adora2a, Penk, Gpr6, Sp9*) were not found to be differentially expressed in the *Shank3*^Δ11/Δ11^ animals (**Table S2**). Moreover, using Single Molecule RNA Fluorescence In Situ Hybridization with *Drd1* and *Drd2* RNA probes, we observed no significant difference in the number of D1 and D2 striatal neurons between the *Shank3*^Δ11/Δ11^ and *Shank3*^+/+^ mice (**Fig. S10**). Altogether, these results indicate that the deletion of exon 11 of *Shank3* leads to differential transcriptional alterations in the D1- and D2-MSN, without major difference in the number of these types of neurons throughout the whole striatum.

**Figure 3:**
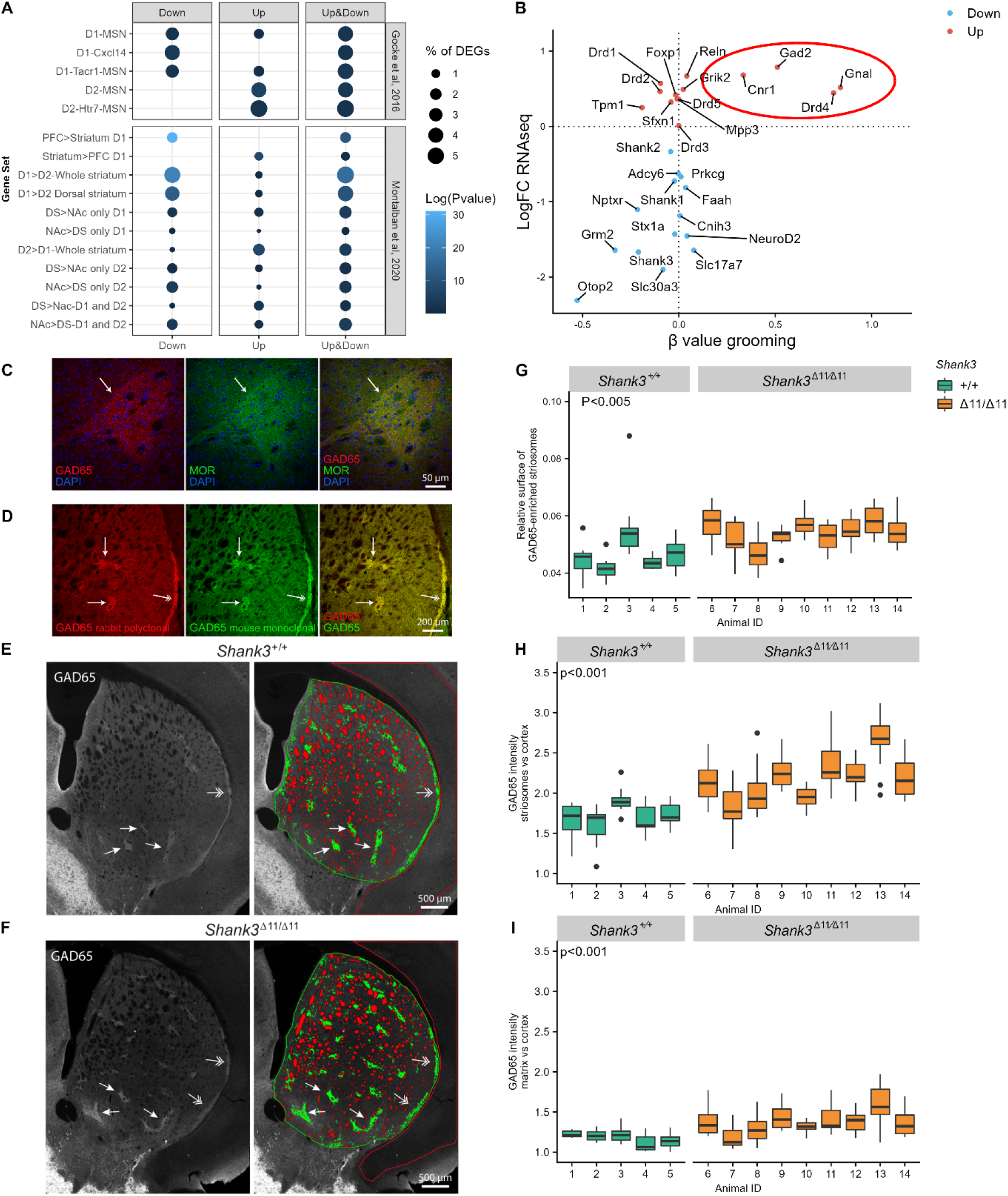
Altered transcriptome of D1- and D2-MSN, over-expression of GAD65, and modification of the striosome/matrix balance in the striatum of *Shank3*^Δ11/Δ11^ mice. (A) Enrichment of RNAseq DEGs in the striatum of *Shank3*^Δ11/Δ11^ mice compared to cell-specific gene clusters suggested by Gokce et al. (2016) and Montalban et al. (2020). Genes under-expressed in *Shank3*^Δ11/Δ11^ striatum are enriched in D1-MSN clusters while genes over-expressed in Shank3^Δ11/Δ11^ striatum are enriched in D2-MSN clusters. Ratio genes correspond to the proportion of DEGs enriched into the cluster. Over-representation analyses were performed with Fisher’s hypergeometric tests and p-values were adjusted for multiple testing with the Benjamini-Hochberg procedure within each category of DEGs. (B) Comparison of the log of the fold change of gene expression of selected genes in *Shank3*^+/+^ vs *Shank3*^Δ11/Δ11^ mice (LogFC RNAseq, Y axis) and the βvalues (slope of the linear regression) for low self-grooming vs high self-grooming (β value grooming, X axis). (C-D) Confocal images of striatum coronal sections of one-year old *Shank3*^+/+^ male mice. (C) GAD65 immunoreactivity (red) is enriched in striatal microzones identified as striosomes by immunostaining for μ-opioid receptor (MOR, in green), a canonical marker of striosomes. (D) Increased GAD65 immunoreactivity in striosomes (arrows) and subcallosal streak (two-headed arrows) compared to surrounding matrix is observed with two different antibodies: the rabbit polyclonal antibody (Invitrogen PA5-77983 in red) used in C, E, F and a mouse monoclonal antibody (Millipore MAB351, in green). (E-F) Distribution and quantification of GAD65 immunoreactivity in dorsal striatum sections of a *Shank3*+/+. (E) and a *Shank3*^Δ11/Δ11^ (F) mouse (images generated by stitching multiple maximum-intensity projected z-stacks). The arrows point to striosomes and the arrowheads to subcallosal streak. In the dorsal striatum ROI (surrounded in green), the unlabelled myelinated fibres are coloured in red, the regions of higher expression of GAD65 (striosomes) are coloured in green, and the regions of lower expression of GAD65 (matrix) are uncoloured. The cortex ROI used as a reference is surrounded in red. Note that the acquisition parameters were adjusted for each brain hemi-section in order to have no saturating signal for GAD65 in the striatal ROI. Increased intensity in the striatum thus leads to a staining which appears weaker in the cortex. (G-I) Comparison of GAD65 immunoreactivity in the striosome and matrix compartments of the dorsal striatum in 5 *Shank3*^+/+^ (green) and 9 *Shank3*^Δ11/Δ11^ (orange) one-year old male mice. (G) Relative surface of the GAD65-enriched striosome compartment (surface of striosomes / surface of (striosomes + matrix). (H) Relative GAD65 labelling intensity in the striosomal compartment of the striatum compared to cortex. (I) Relative GAD65 labelling intensity in the matrix compartment of the striatum compared to cortex. Data, generated from analysis of 9 to 14 images per animal, are presented as box-plots (median, first, and third quartiles).

### Abnormal expression of striosome markers

Using RT-PCR, we investigated the correlation between the mRNA levels in the striatum of DEGs identified by RNAseq analysis and other genes of interest (*Shank1, Shank2, and Drds*) and the level of self-grooming observed in seven 12 months-old *Shank3*^Δ11/Δ11^ mice (**Fig S11**). We found that excessive grooming correlated with higher levels of transcripts for four of the DEGs identified by RNAseq or RT-PCR: *Cnr1, Gnal, Gad2*, and *Drd4* (**Fig. 3B**). *Cnr1* encodes the cannabinoid receptor 1; *Gnal* encodes a stimulatory G-alpha subunit of a heterotrimeric G-protein (Gαolf); *Gad2* codes for GAD65, one of the two enzymes that convert glutamate into γ-aminobutyric acid (GABA), and *Drd4* encodes the dopamine receptor 4. Interestingly, these proteins have been reported to be heterogeneously distributed in the dorsal striatum, being more abundant in the spatial microzones known as striosomes or patches than in the much larger matrix surrounding them, especially *Gad2/*GAD65 ^23,24^. Therefore, given the function of GAD65 (i.e. converting glutamate into GABA, an inhibitory transmitter), we studied its distribution in the dorsal striatum, using immunofluorescence on brain sections (**Fig. 3C-I**). Using two different anti-GAD65 antibodies, we found that GAD65 is enriched in striosomes, identified by co-labelling of µ-opioid receptor (MOR), the most widely used marker for striosomes of the rostral striatum ^25^ (**Fig. 3C, D**). GAD65 can thus be considered as a marker of the striosomal compartment in mice. Quantitative analysis revealed that the relative surface occupied by the GAD65-enriched striosomes in the dorsal striatum is increased in the *Shank3*^Δ11/Δ11^ mice compared to the *Shank3*^+/+^ mice (p = 0.0018, **Fig. 3E-G**). Moreover, in agreement with the overexpression of *Gad2* in the striatum but not in the cortex (table S2) of *Shank3*^Δ11/Δ11^ mice, GAD65 labelling intensity relative to cortex was increased in the whole *Shank3*^Δ11/Δ11^ striatum, but at much higher levels in the striosomes (p= 6.5 10^−5^, **Fig. 3G**) than in the matrix (p= 2.9 10^−5^, **Fig. 3H and Fig. S12A, B**). SHANK3 expression, observed using a specific antibody directed against the C-terminal region present in the majority of SHANK3 isoforms, is comparable in the striosomal and matrix compartments of the striatum in both *Shank3*^+/+^ and *Shank3*^Δ11/Δ11^ mice (**Fig. S12A, C**). Finally, we investigated GAD65 distribution in the striatum of another *Shank3* mouse model, lacking all SHANK3 isoforms ^9^. These 20-28 weeks-old *Shank3*^Δ4-22/Δ4-22^ male mice were also used to validate the specificity of the anti-SHANK3 antibody (Fig. S12). We confirmed enlargement and GAD65 over-expression in the striosomal compartment. However, in the absence of all SHANK3 isoforms, GAD65 is also markedly over-expressed in the matrix (**Fig. S12**).

## Discussion

The present study highlighted robust behavioural deficits in *Shank3*^Δ11/Δ11^ mutant mice, more specifically a reduced activity in the open field, atypical social behaviour in males interacting with oestrous females, and increased self-grooming compared to wild-type littermates. The majority of *Shank3*^Δ11/Δ11^ mutant mice displayed a tendency to a worsening of the self-grooming phenotype with increasing age for mice showing an early emergence of this trait. At 12 months of age, the striatum had the most impacted transcriptomic profile compared to other brain regions. Further molecular characterizations pointed towards possible imbalances between the striosome and matrix compartments in the dorsal striatum.

### Behavioural profiling of the Shank3^Δ11/Δ11^ mutant mice

#### Atypical social interaction

Results from the literature have highlighted either reduced duration of social contact and reduced number of USVs emitted [e.g., *Shank3* ^*Δex4-2* 26^] or no significant difference in socio-sexual behaviours [e.g., *Shank3*^Q321R 27^]. Here, we observed limited difference in social interaction with an increase in the time spent in contact for *Shank3*^Δ11/Δ11^ males interacting with an oestrous female (observed at 3 and 8 months of age), and more specifically in nose-to-nose contacts, as already highlighted in previous studies [*Shank3*Δ^ex14−16^cKO ^27^; *Shank3*^Δ11/Δ11^ females ^28^]. Testing the specificity of this type of contact would help to understand its significance in mouse social communication. Further experiments should be designed to test whether this specific behaviour is associated with a deficit in social reward processing [as suggested for *Shank2*^Δ6-7/Δ6-7^ mice ^29^]. Indeed, several studies have shown that SHANK3 controls maturation of social circuits in the ventral tegmental area (VTA) ^30,31^ and synaptic strength of D2-MSN in the striatum ^13,18^.

#### Excessive grooming

Excessive self-grooming has been observed in the majority of *Shank3* mutant mice over their lifetime and therefore represents the most robust behavioural phenotype in all models. Extreme self-grooming is most likely not due to increased skin sensitivity since rescuing normal tactile reactivity in *Shank3b*^+/-^ does not improve over-grooming behaviour ^32^. In our study, we also observed an important inter-individual variability in this trait with 30% of the mutant mice that did not seem to display excessive grooming even after one year. Increased self-grooming might reflect a response to stressful conditions, which might vary from one individual to another (difference in susceptibility and/or exposure to stressful conditions). Such increased reactivity to novel or stressful conditions is reminiscent of what is observed in patients and deserves further systematic testing in the different *Shank3* mutant models ^33^.

### From transcriptome to imbalance of striatum compartments and excessive self-grooming

#### Abnormal striatal transcriptome

The massive impact of the *Shank3*^Δ11^ mutation on the transcriptome profile of the striatum suggests that this region could be the centrepiece of the behavioural deficits observed in *Shank3*^Δ11/Δ11^ mutant mice. Several studies have already reported functional alterations of the striatum and more specifically of the MSN in the diverse *Shank3* mutant strains ^10,13,15,18,30,31^. Our report of an alteration of the striatal transcriptome could be due to a direct role of the SHANK3 protein in gene transcription. Indeed, a previous study, using transfected cells expressing EGFP-SHANK3 fusion proteins, has reported the presence of SHANK3 in the nucleus, especially the isoform SHANK3b ^7^, which is disrupted in our model. Using immunohistofluorescence, we did not observe SHANK3 in the nucleus of the striatal cells in *Shank3*^+/+^ adult mice, but the epitope recognized by the antibody we used is absent from the SHANK3b isoform. Alternatively, the abnormal striatal transcriptome profile could be the consequence of the alteration of the glutamatergic synapses that interfere with signal transduction, such as G protein ^34^ and mTOR ^35^ signalling, and then with gene transcription or mRNA stability. We found that several transcripts encoding proteins involved in glutamatergic synaptic transmission were under-expressed in *Shank3*^Δ11/Δ11^ mice. For example, we observed a decrease of *Grm2*, encoding mGluR2, a G-protein-coupled glutamate receptor which interacts with the proline-rich domain of SHANK3 and which has been found decreased in a valproate-induced rat model of autism ^36^. We also observed significant under-expression of *Dlgap1*, coding for DAP1/GKAP. This protein connects SHANK3 to the scaffolding protein PSD-95 ^37^ which recruits AMPAR and NMDAR. In contrast, we did not observe diminution of transcript levels for several proteins (GluA, GluN or mGluR5) previously shown as decreased in the synaptosomes or PSD fractions of *Shank3* mutant mice compared to wild type. A possible explanation is that the absence of SHANK3 decreases the translation or the recruitment of these proteins at the synapse or increases their degradation without affecting gene transcription and mRNA degradation. Interestingly, we observed a sexual dimorphism in behaviour in mice, especially for cohort 1, that is not reported in patients (Leblond et al., 2014). This aspect could be explained by the impact of oestradiol on striatal function and behaviour in mice ^38^, as well as by differential gene expression between males and females after an acute stress in mice as, for instance, in the CA3 subregion of hippocampus ^39^. Analysing the brain transcriptome of *Shank3*^+/+^ and *Shank3*^Δ11/Δ11^ females should shed light on the differential effect of the lack of *Shank3* between sexes.

#### A link between abnormal gene transcription, striatum organisation and behavioural impairment

We observed that the expression of *Gad2, Cnr1, Gnal* and *Drd4* was positively correlated with excessive self-grooming. Remarkably, three of these genes have been previously associated with self-grooming, but not always in the same direction. In contrast to our study, *Cnr1* deletions (as well as CB1R antagonists) have been reported to increase self-grooming behaviour ^40,41^. The overexpression of *Cnr1* in *Shank3*^Δ11/Δ11^ mice might therefore result from a compensatory upregulation in response to the alterations of the endocannabinoid system which has been previously reported in SHANK3-deficient mice ^18,42^, in valproic acid induced rat models of autism and also in some autistic individuals ^43^. Consistent with our findings, *Gnal* haploinsufficiency is associated with a reduction of self-grooming behaviour in a mouse model of dystonia ^44^. Links between *Gad2* and *Shank3* were previously reported with an increase of GAD65 immunoreactivity in the *Shank3*ß^-/-^ mice ^40^ and a decrease of *Gad2* transcript in the striatum of a *Shank3*-overexpressing model ^45^. GAD65 is associated with different disorders related to anxiety, such as obsessive-compulsive disorder (OCD), panic disorder or generalized anxiety disorder ^46^. To our knowledge, there is no direct evidence in the literature for an association between *Drd4* expression and grooming behaviour. However, *Drd4* is a candidate gene for OCD and panic disorder ^47^. It would thus be interesting in future studies to investigate this potential link.

Interestingly, these four genes have also been reported to be enriched in striosomes ^23,24^. These microzones of the dorsal striatum contain early born MSN and are embedded into the much larger surrounding matrix including late-born MSN ^48^. The two compartments, initially distinguished by differential expression of numerous molecular markers [for a review see ^49^], also differ in their input and output connectivity and electrophysiological characteristics ^50^, such as D1-MSN excitability ^51^, rates of dopamine release ^52^, and response to chronic stress ^53^. Imbalances in striosome to matrix activity have been suggested in several movement disorders [Huntington’s disease, Parkinson’s disease, dyskinesia, dystonia, for a review see ^49^], psychostimulant-induced motor stereotypies ^54^, and more recently in anxiety disorder ^46^. Corticostriatal path targeting striosomes also control decision-making under cost-benefit conflict ^55,56^ and habit formation ^57^. Our finding that four striosome markers had increased expression associated with excessive self-grooming in *Shank3*^Δ11/Δ11^ mice prompted us to explore the compartmental architecture of the striatum in these mice. This investigation was also motivated by previous studies that have shown that enhanced striosomal activation is highly correlated with increased repetitive behaviours, including self-grooming in both non-human primates and rodents [for review see ^12^]. Studies by Kuo and Liu ^58,59^ have recently suggested that aberrant striatal compartmentation may be involved in autism. Furthermore, through the investigation of protein-protein interactions, a role for the striosomes in anxiety disorder such as social anxiety has been recently proposed ^46^.

We first confirmed that GAD65, which had been reported to be a striosome marker in primates, but not in rats ^24^ was indeed a striosome marker in the mouse. We then found that the GAD65-enriched striosomal compartment is enlarged in *Shank3*^Δ11/Δ11^ mice compared to *Shank3*^*+/+*^ mice and that striatal GAD65 overexpression in the absence of the major SHANK3 isoforms is much more pronounced in the striosomes than in the matrix. Whereas the majority of GABA in brain is produced by the cytosolic isoform of glutamate decarboxylase (GAD67, encoded by *Gad1*), GAD65, which is anchored to the membrane of synaptic vesicles, can supply GABA in situations of high demand ^60^, such as stress or fear, and for fine tuning of inhibitory transmission. It is thus possible that an alteration of striosomes in *Shank3*^Δ11/Δ11^ mice is responsible for some of the behavioural features such as excessive self-grooming. *A model for striosomes/matrix imbalance as a possible cause of excessive grooming in the Shank3*^Δ11/Δ11^*mice:* It is well established that the striatum and the dopamine-containing nigrostriatal tract control self-grooming behaviour ^12^, and that striosomal MSN are the predominant striatal population innervating dopamine neurons of the nigrostriatal tract ^61^. An imbalance between the striosome and the matrix compartments may then explain, via a modification of dopamine signalling, the excessive grooming of *Shank3*^Δ11/Δ11^ mice (**Fig. 4**). Indeed, in contrast to MSN located in the matrix of the striatum, striosomal MSN projects directly to the dopamine-producing neurons in the SNpc ^49,62^. Then, SNpc dopamine neurons project back to the entire dorsal striatum. An increased activity of GAD65 in striosomes, in a situation of stress for example, would lead to an inhibition of dopamine release by the SNpc and thus to a reduced dopamine modulation of both the direct and indirect pathways in the dorsal striatum. Because dopamine activation has opposite effects on D1- and D2-MSN by increasing and decreasing cell excitability, respectively, a decreased dopamine release by SNpc is expected to result in diminishing D1-MSN activity and boosting MSN-D2 activity. The differential enrichment in up- and down-regulated DEGs in D2-MSN and D1-MSN, respectively, could be related to previous observation of a distinct impairment of LTD in D2-MSNs, but not in D1-MSN ^18^ in *Shank3* mutant mice. Importantly, striosome cells are generated before matrix cells during development ^48^ and SHANK3 seems to be involved in the early stages of neuronal development ^15,63^. Therefore, the enlargement of the total relative surface of the striosomal compartment observed first in the *Shank3*^Δ11/Δ11^ and then in the *Shank3*^Δ4-22/Δ4-22^ mice may be due to a default in the compartmentation of the striatum during early development.

**Figure 4:**
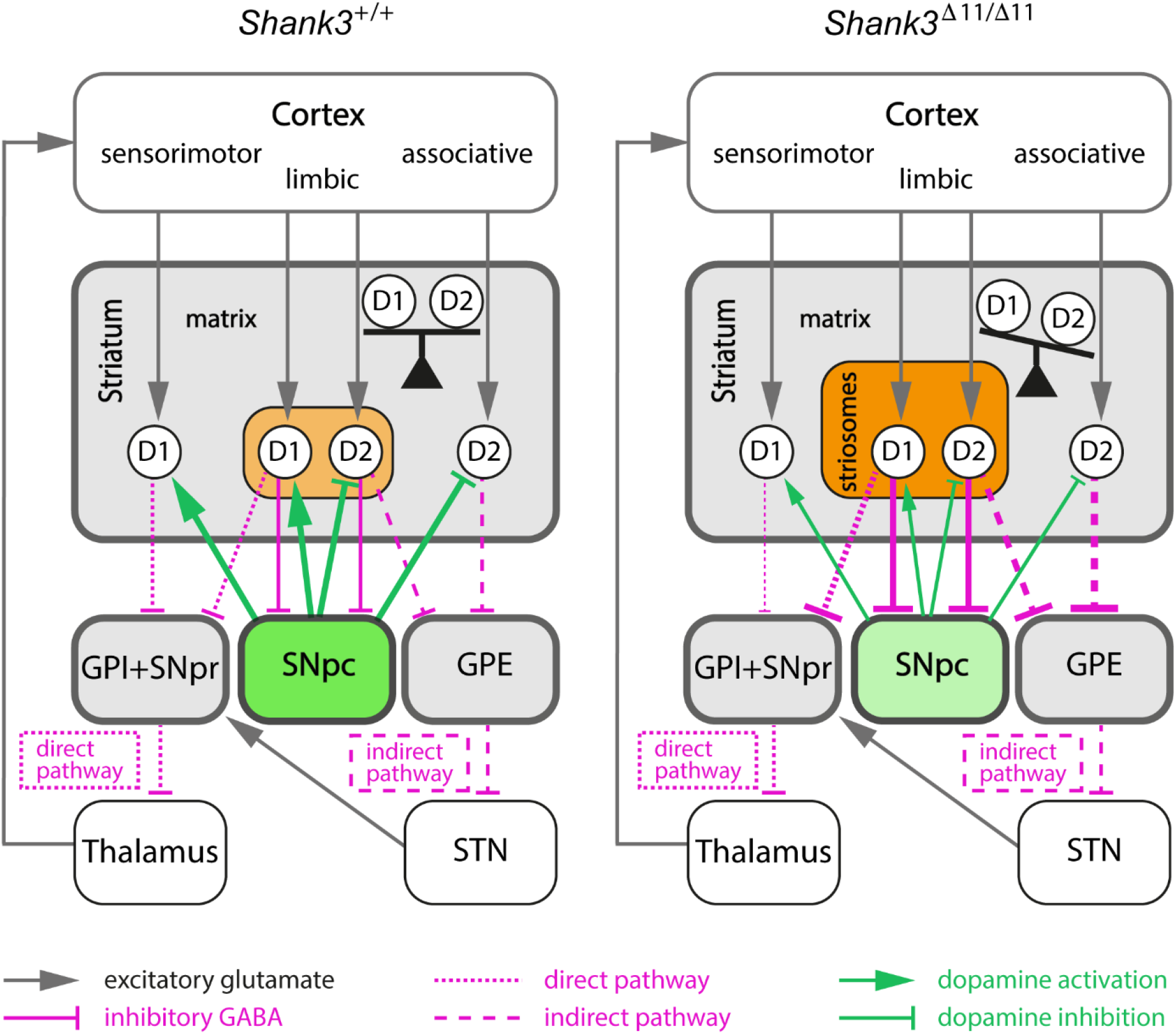
Proposed model linking increased activity in the striosomal pathway and imbalance between the direct and indirect pathways in the *Shank3*^Δ11/Δ11^ mice. The modifications of connection strength (line thickness) induced by striosomal compartment alterations in *Shank3*^Δ11/Δ11^ mice are only represented between the striatum and its output target areas (colored boxes). Increased activity of the striosomal pathway in *Shank3*^Δ11/Δ11^ mice, due to both larger size and increased GAD65 activity of the striosomal compartment (light/dark orange) enhances GABAergic inhibition of the dopamine-producing neurons of SNpc (green). This inhibition of the SNpc neurons (light green) results in an imbalance between the direct and indirect pathways by decreasing both the activation of D1-MSN (D1) and the inhibition of D2-MSN (D2). GPI, internal globus pallidus; GPE, external globus pallidus; SNpr, substantia nigra pars reticulata; SNpc, substantia nigra pars compacta; STN, subthalamic nucleus.

### Conclusions and perspectives

Altogether, the present study describes the effects over time of the *Shank3*^Δ11^ mutation on mouse social and repetitive behaviours and suggests a critical role of the striatum in excessive self-grooming, probably through imbalances in two intertwined systems, the striosome/matrix compartments and the direct/indirect pathways. Our results suggest that SHANK3 deficiency/Phelan McDermid syndrome and possibly to some extent autism could integrate the growing list of neurological conditions implicating an imbalance in the striosome/matrix compartmentation of the dorsal striatum. Future studies in mice, but also in other species such as rat ^64^ or non-human primates ^65,66^, are necessary to understand the mechanisms at the origin of the striosome/matrix imbalance and consider the possibility of its reversibility.

## Material and methods

### Animals and cohorts

*Shank3*^Δ11/Δ11^ mice were generated by Genoway (Lyon, FRANCE) using 129S1/SVImJ ES cells. The *Shank3* mutation, deletion of exon 11, was then transferred onto a C57BL/6J background with more than 15 backcrosses ^19^. In this model, some protein isoforms are still expressed ^19^. Food and water were provided *ad libitum*. The experimenters were blind to the genotype of the tested animals for data collection and analyses. We bred *Shank3*^Δ11/+^ males and females to obtain *Shank3*^+/+^, *Shank3*^+/Δ11^ and *Shank3*^Δ11/Δ11^ littermates. For behavioural tests, we used three cohorts (including both males and females) that we analysed separately. See **Supplementary Methods** for details on the cohorts. *Shank3*^Δ4-22/Δ4-22^ mice ^9^, with a C57BL/6N background, were obtained from The Jackson Laboratory Repository. Ethical statement

All procedures were validated by the ethical committees of the Institut Pasteur (CEEA n°89) and of the Sorbonne Université (CEEA n°5) and were conducted under the approval of the Ministère de l’Enseignement Supérieur, de la Recherche et de l’Innovation under the reference APAFIS#7706-20 161 12317252460 v2 and reference APAFIS#3234-2015121619347680 v7, respectively.

### Behavioural tests

Dark/light test, Elevated Plus-maze test, locomotion and exploratory test in the open field, analysis of stereotyped behaviour, occupant/new-comer social test with ultrasonic vocalisation recording, male behaviour in presence of an oestrous female, three-chambered social test and starmaze test were used to investigate the mice’s behaviours. The protocols of these tests are detailed in the “behavioural tests” section of the **Supplementary Methods**.

### Behavioural statistics

All group data are represented as mean ± standard error of the mean (s.e.m.), as well as the individual points. All statistics were performed with R software (R Core Team,R Foundation for Statistical Computing, 2020). In all behavioural tests, *Shank3*^+/Δ11^ mice were not significantly different from *Shank3*^+/+^ unless otherwise specified (see **Table S1**). Given the limited sample size and the non-normality of the data, comparisons between genotypes were performed using non-parametric Mann-Whitney U-tests. We used the non-parametric Friedman test to examine the effect of age within each genotype. Between age points, post-hoc tests were performed using paired Wilcoxon signed-rank tests. If required, a Bonferroni correction for multiple testing was performed. Differences between groups were considered significant when p<0.05.

### Transcriptome analysis

Tissue collection, total RNA extraction and sequencing, mapping and reference genome, calling variants from the RNAseq data and differential expression analysis are detailed in the “Transcriptome analysis” section of the **Supplementary Methods**.

### Gene set and protein-protein interaction network analysis

Gene set over-representation analyses were performed based on Ensemble of Gene Set Enrichment Analyses (EGSEA) without considering the genes around the *Shank3* gene (see details in the **Supplementary Methods)**. Protein-protein interaction (PPI) networks were generated using the DEGs for the four brain structures independently. A randomised protein interaction network was created using all DEGs to determine the probability of finding a network. Software, packages and databases are detailed in the “Gene set and protein-protein interaction network analysis” section of the

## Supplementary Methods

### Quantitative RT-PCR (q-RT-PCR) and RNA and protein fluorescent detection on striatum sections

We selected several genes of interest by cross checking the DEGs, ORA (gene ontology – GO –, Kyoto encyclopaedia of genes and genomes – KEGG– and Pathway), and data from the literature. To validate these genes, we used q-RT-PCR, either with the droplet digital Polymerase Chain Reaction (dd-PCR) or with the real time quantitative PCR (q-PCR) technology. These procedures are detailed in the “Quantitative RT-PCR” section of the **Supplementary Methods**. All statistical analyses were performed using R software (R Core Team, R Foundation for Statistical Computing, 2020). The comparison between genotypes was performed using Mann-Whitney U-tests. Single-molecule Fluorescent In Situ Hybridization (smFISH) experiment to detect *Drd1* and *Drd2* RNAs and immunofluorescence experiment and image analysis to quantify GAD65 immunoreactivity in the dorsal striatum are detailed in the “Single-molecule Fluorescent In Situ Hybridization and immunofluorescence experiments on brain sections” and “Quantitative analysis of GAD65 immunoreactivity” section of the **Supplementary Methods**.

## Acknowledgments

The authors thank the members of the Génétique Humaine et Fonctions Cognitives lab for helpful discussions, especially Jean-Pierre Bourgeois for his advice and expertise, Thomas Rolland for his assistance with data analysis and Nathalie Lemière for her assistance with sample collection. The authors thank Jean-Antoine Girault, Véronique Brault, and Sonia Garel for their advice and expertise.

## Detailed author contributions

ATF, ABi, EV, EE and TB conceived the experiments. ATF and EE designed and acquired behavioural data from cohorts 1 and 2. ATF extracted and dissected brains, extracted RNA from cohorts 1 and 2. ATF and FM designed and ATF acquired data from smFISH. ATF performed analysis and statistics of behaviour, PPI and smFISH. ABo, BFi and JFD performed the RNA sequencing. ABi and ATF performed the analyses of ORA. ATF, BFo and AL performed and analysed the results of quantitative RT-PCR. EV performed the immunofluorescence experiments, with the help of SC for brain collection. FdC performed quantitative image analysis. SC and AMLS genotyped the mice. JS, CR and LRR conducted the experiments on cohort 3. TMB generated the *Shank3*^Δ11/Δ11^ mouse model. ATF, ABi, EV, BF, FdC, EE and TB wrote the manuscript.

## Data and materials availability

All data and material are available upon request to the corresponding authors.

