## Supplementary data for "Excessive self-grooming, gene dysregulation and imbalance between the striosome and matrix compartments in the striatum of *Shank3* mutant mice"

Supplementary Figures 3
- Supplementary Figure S1 3
- Supplementary Figure S2 4
- Supplementary Figure S3 5
- Supplementary Figure S4 6
- Supplementary Figure S5 7
- Supplementary Figure S6 8
- Supplementary Figure S7 10
- Supplementary Figure S8 11
- Supplementary Figure S9 12
- Supplementary Figure S10 13
- Supplementary Figure S11 14
- Supplementary Figure S12 15
- Supplementary Figure S13 17
- Supplementary Figure S14 18

Supplementary Material & Methods 19
- Animals and *Shank3*$^{\Delta 11}$ cohorts 19
- Behavioural tests 19
- Transcriptome analysis 23
- Gene set and protein-protein interaction network analysis 26
- Quantitative RT-PCR 27
- Single-molecule Fluorescent In Situ Hybridisation and immunofluorescence experiments on brain sections 27

### Supplementary Figures

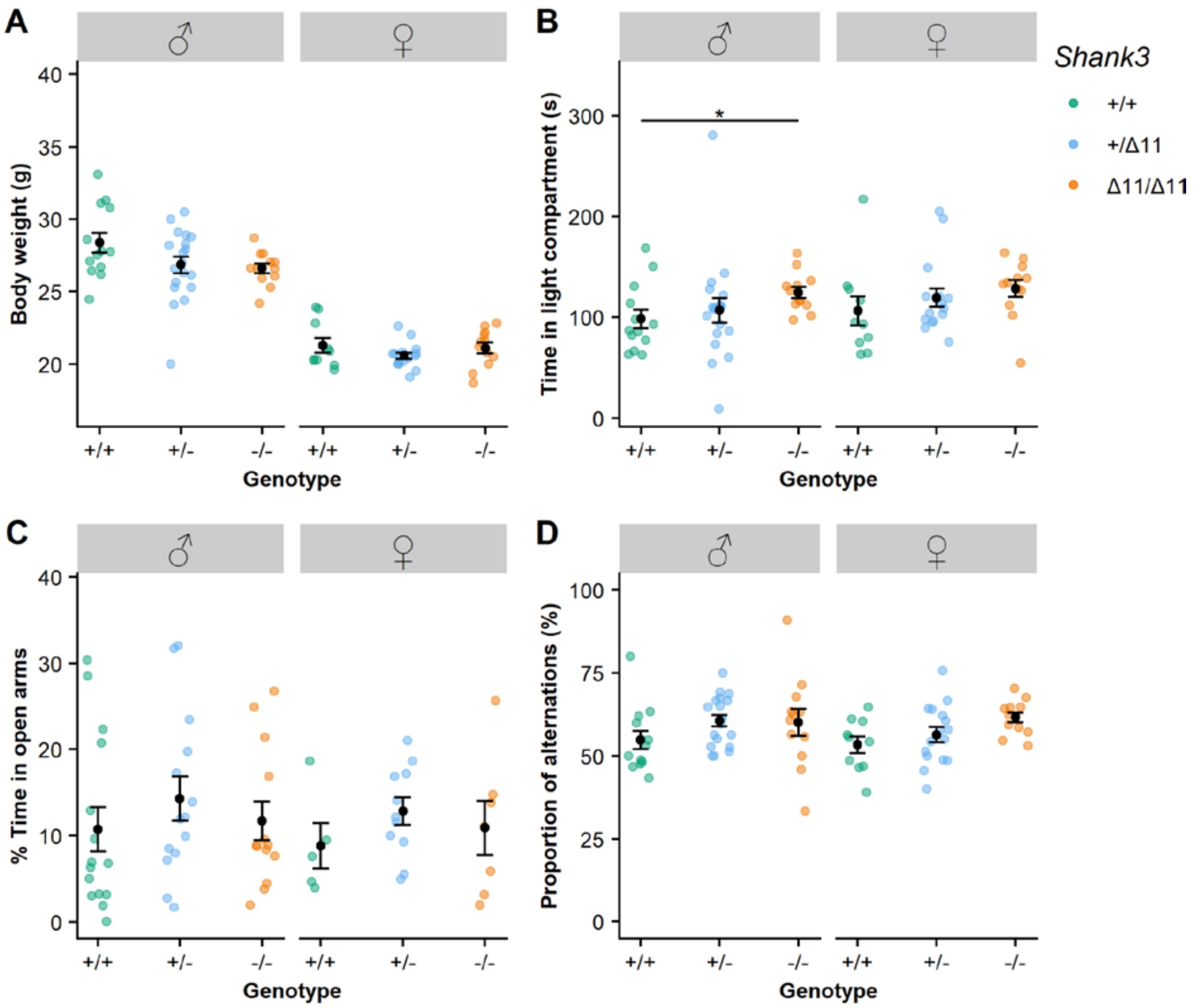


#### Supplementary Figure S1

Three month-old *Shank3*$^{\Delta11/\Delta11}$ mice display subtle genotype-related differences with their *Shank3*$^{+/+}$ and *Shank3*$^{+/\Delta11}$ littermates in weight, motor coordination, anxiety and working memory. (A) Body weight of male (left panel) and female (right panel) of Cohort 1. (B) Time spent in the light compartment in the dark/light test during 5 min for Cohort 1. (C) Proportion of time spent in the open arms in the elevated plus maze during 10 min for Cohort 2. (D) Proportion of alternations between the three arms of a Y-maze for Cohort 1. Mann–Whitney U test with Bonferroni correction for multiple testing: *: corrected p-value <0.05; data are presented as mean ± s.e.m. (black squares and circles); 12–18 mice per group.

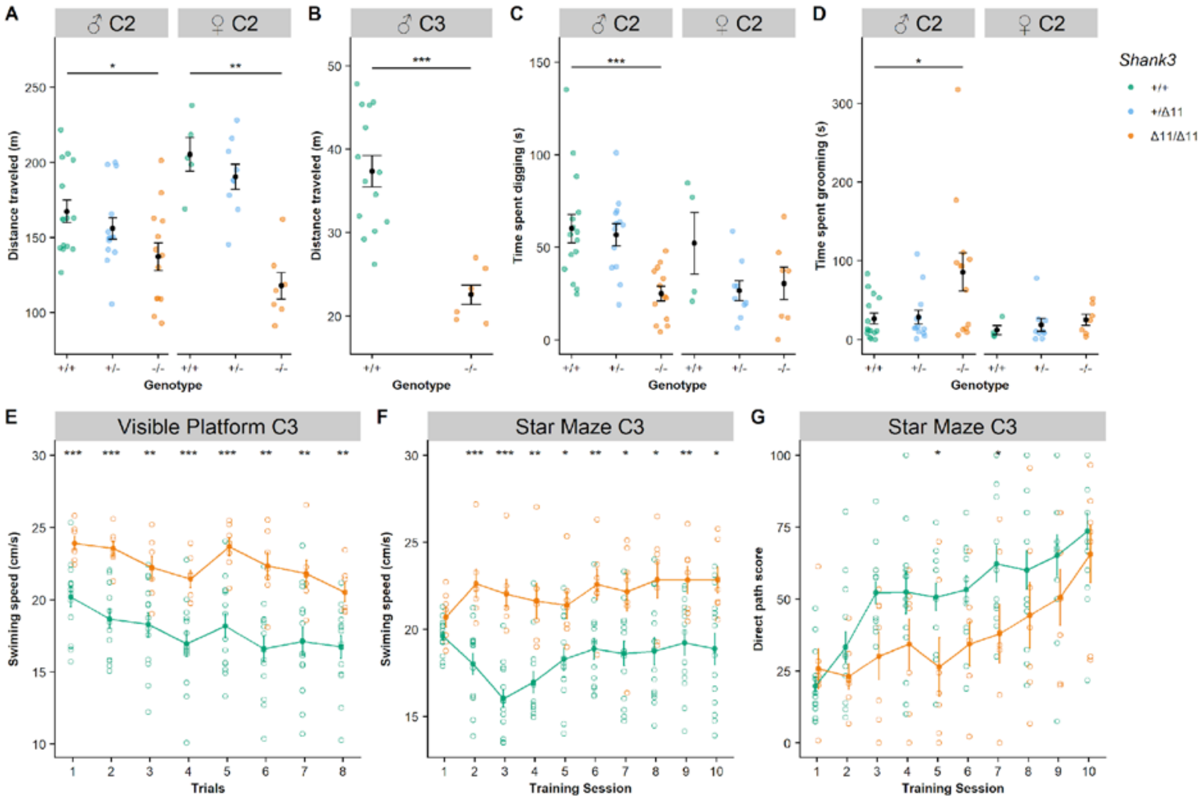


### Supplementary Figure S2

*Shank3*$^{\Delta11/\Delta11}$ mice of Cohorts 2 (C2) and 3 (C3) at three months of age display atypical activity and exploration and increased stereotyped behaviour. (A, B) Total distance travelled during free exploration of an open-field for Cohort 2 (C2) during 30 min (A) and Cohort 3 (C3) during 10min (B). (C) Total time spent digging, i.e. moving the bedding with front and/or hind legs, during 10 min observation in a test cage (after 10 min habituation). (D) Total time spent self-grooming during 10 min observation in a test cage with fresh bedding (after 10 min habituation). (E) Swimming speed in a Morris water maze with a visible platform. (F) Swimming speed in a starmaze. (G) Direct-path score in the Starmaze test. Mann–Whitney U test, with Bonferroni correction for multiple testing (in black): *corrected p-value <0.05, **corrected p-value <0.01, ***corrected p-value <0.001; data are presented as mean ± s.e.m.; cohort 2: 8–22 mice per group; cohort 3: 7–14 mice per group.

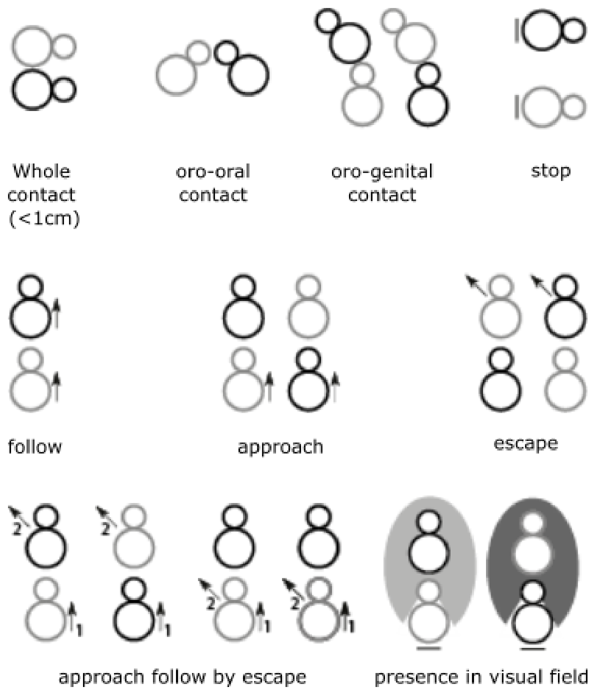


**Supplementary Figure S3**

Behavioural events detected by Mice Profiler during social interactions between the occupant (light grey) and the new-comer (dark grey). Arrow defines the movement of one of the animals.

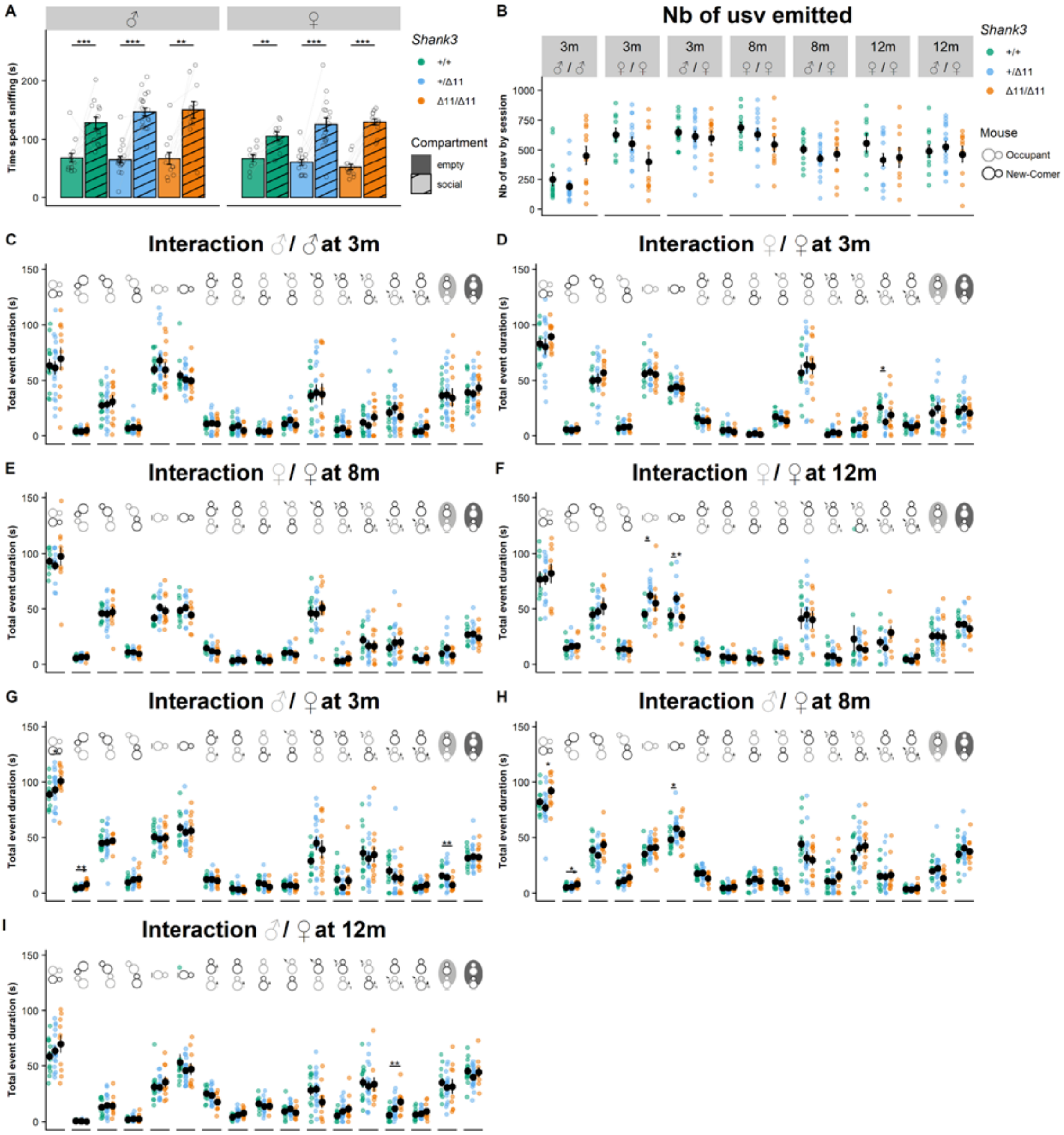


**Supplementary Figure S4**

*Shank3*$^{\Delta11/\Delta11}$ mice display limited genotype-related differences in social events at three, eight and twelve months of age. (A) Social preference measured in time spent sniffing during the first phase of a three-chambered test. (B) Number of ultrasonic vocalisations (USVs) emitted during 4-min dyadic social interaction. (C – I) Time spent in the different types of social events during 4-min male/male (C), male/female (E, G, I) and female/female (D, F, H) interactions. In all free moving tests, the tested mouse (the occupant) is indicated in light grey and interacts with a wild-type C57BL/6J mouse (the new-comer) indicated in dark grey. Mann–Whitney U test, with Bonferroni correction for multiple testing (in black): *corrected p-value <0.05, **corrected p-value <0.01, ***corrected p-value <0.001; data are presented as mean ± s.e.m.; 12–14 mice per group.

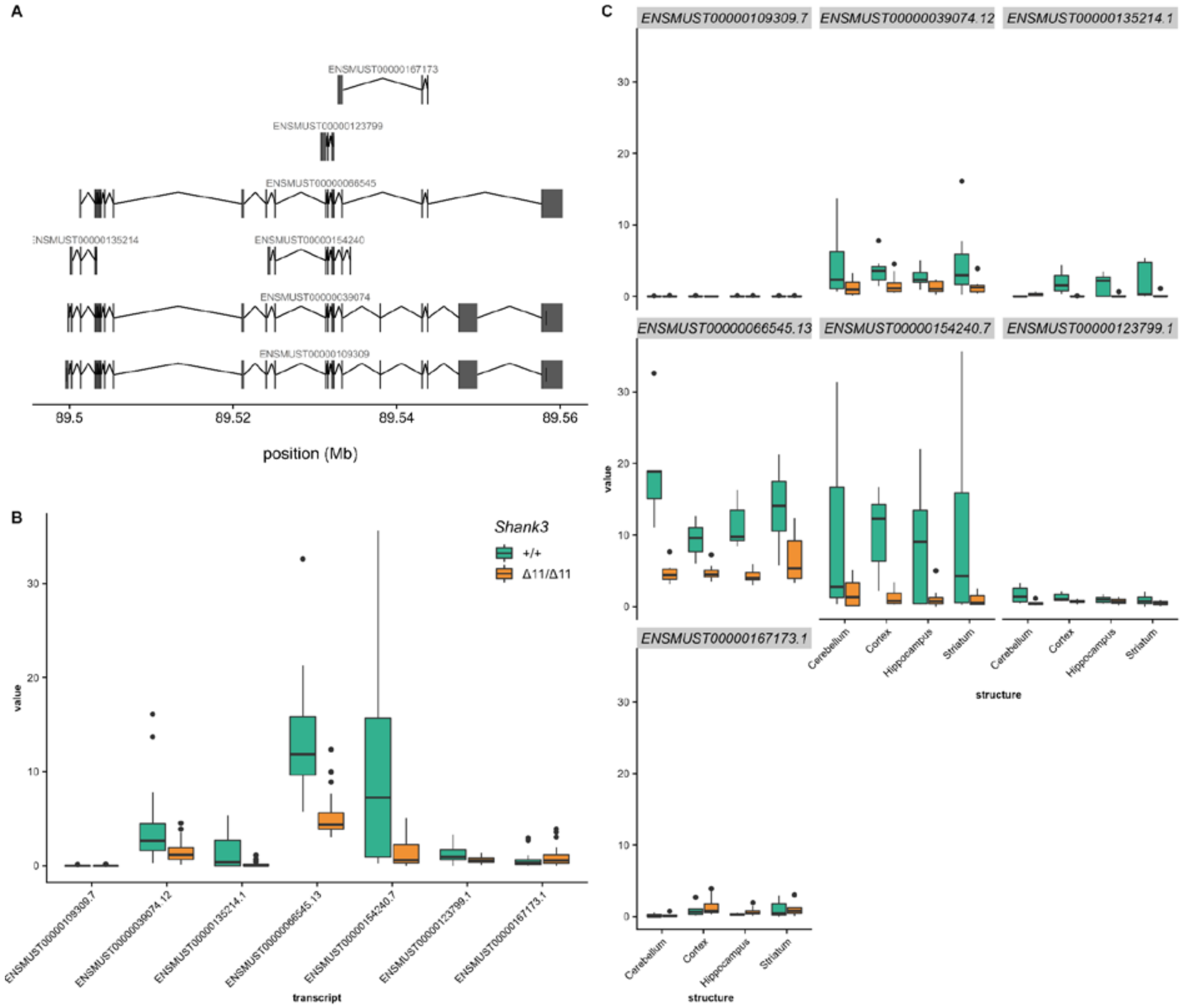


**Supplementary Figure S5**

*Shank3* isoform abundance estimates were computed with salmon (Patro R. et al., 2017) using the *Mus Musculus* GRCm38/mm10 genome and Ensembl transcripts annotations. Isoform abundances are measured in log2 transcripts-per million (TPM). A. *Shank3* Ensembl isoforms B. Box plots of *Shank3* isoform abundances (log2 TPM) across all samples in *Shank3*$^{+/+}$ (green) and *Shank3*$^{\Delta 11/\Delta 11}$ (orange) mice. C. Box plots of *Shank3* isoform abundances (log2 TPM) in *Shank3*$^{+/+}$ (green) and *Shank3*$^{\Delta 11/\Delta 11}$ (orange) mice within each brain structure (cerebellum, cortex, hippocampus and striatum).

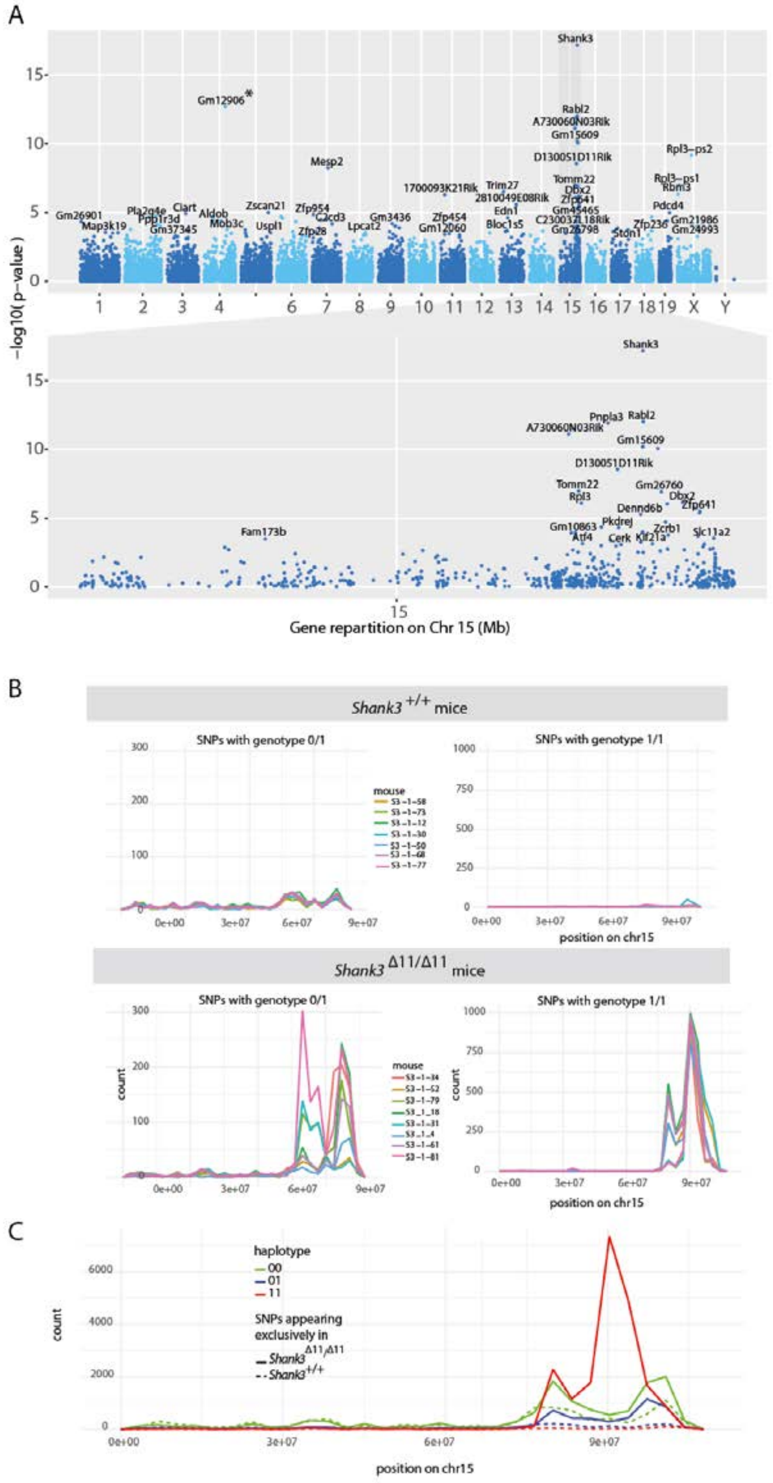

**Supplementary Figure S6**

An increased number of SNP around the *Shank3* locus suggests a construct bias. In the region spanning between 10.4 Mb upstream and 7.5 Mb downstream of *Shank3*, SNPs differing from the reference genome of C57BL/6J mice were found in *Shank3*$^{\Delta11/\Delta11}$ mice, but not in *Shank3*$^{+/+}$ mice. (A) Manhattan plot showing the negative log10-transformed p-value of the comparison (across all brain structures) of gene expression between *Shank3*$^{+/+}$ and *Shank3*$^{\Delta11/\Delta11}$ mice for each read on the whole genome (upper panel) and a close-up view on the chromosome 15 (lower panel). The star (*) indicates the gene *Gm12906*, a pseudogene of the gene *Tomm22* located close to *Shank3* on chromosome 15. (B) Distribution on chromosome 15 of the heterozygous variant genotypes (0/1; right panel) and homozygous variant genotypes (1/1; left panel) that are specific to either *Shank3*$^{+/+}$ (upper panels) or *Shank3*$^{\Delta 11/\Delta 11}$ mice (lower panels). Data are presented as the counts of variants as a function of the genomic position on chromosome 15; each line represents an individual, with samples from all brain regions pooled together. (C) Distribution of the homozygous and heterozygous variants across individuals of the same genotype. Data are presented as the counts of variants as a function of the genomic position on chromosome 15. Homozygous (resp. heterozygous) variant genotypes are shown in red (resp. blue). Plain (resp. dotted) lines represent variants found in *Shank3*$^{\Delta 11/\Delta 11}$ (resp. *Shank3*$^{+/+}$) mice.

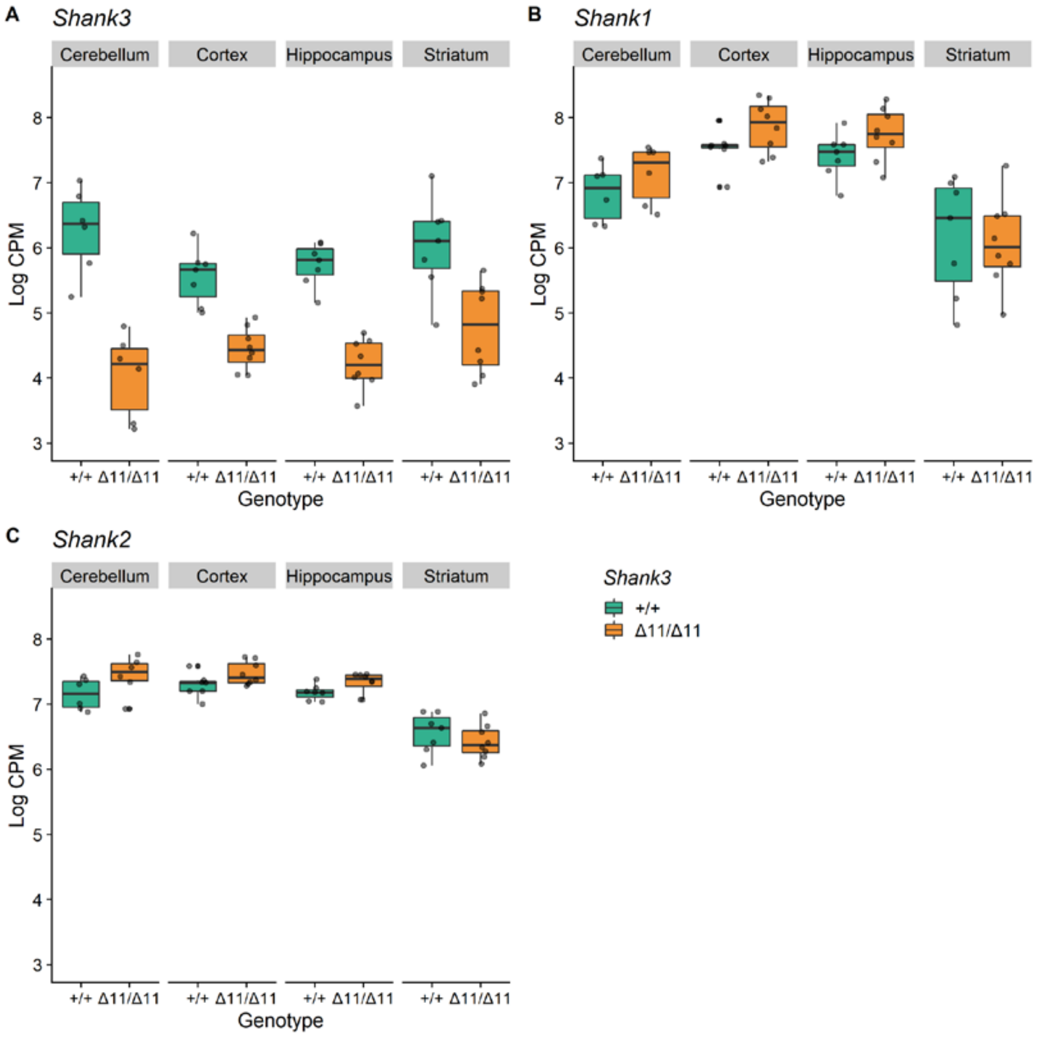


**Supplementary Figure S7**

RNA sequencing-based expression levels of *Shank1*, *Shank2* and *Shank3* in four brain regions. Log counts of reads per million (RPM) for *Shank3* (A), *Shank1* (B) and *Shank2* (C) transcripts in the cerebellum, cortex, hippocampus and striatum for *Shank3*$^{+/+}$ (green) and *Shank3*$^{\Delta11/\Delta11}$ (orange) mice. Data are presented as box-plots (median and first and third quartiles) and sample points (black points).

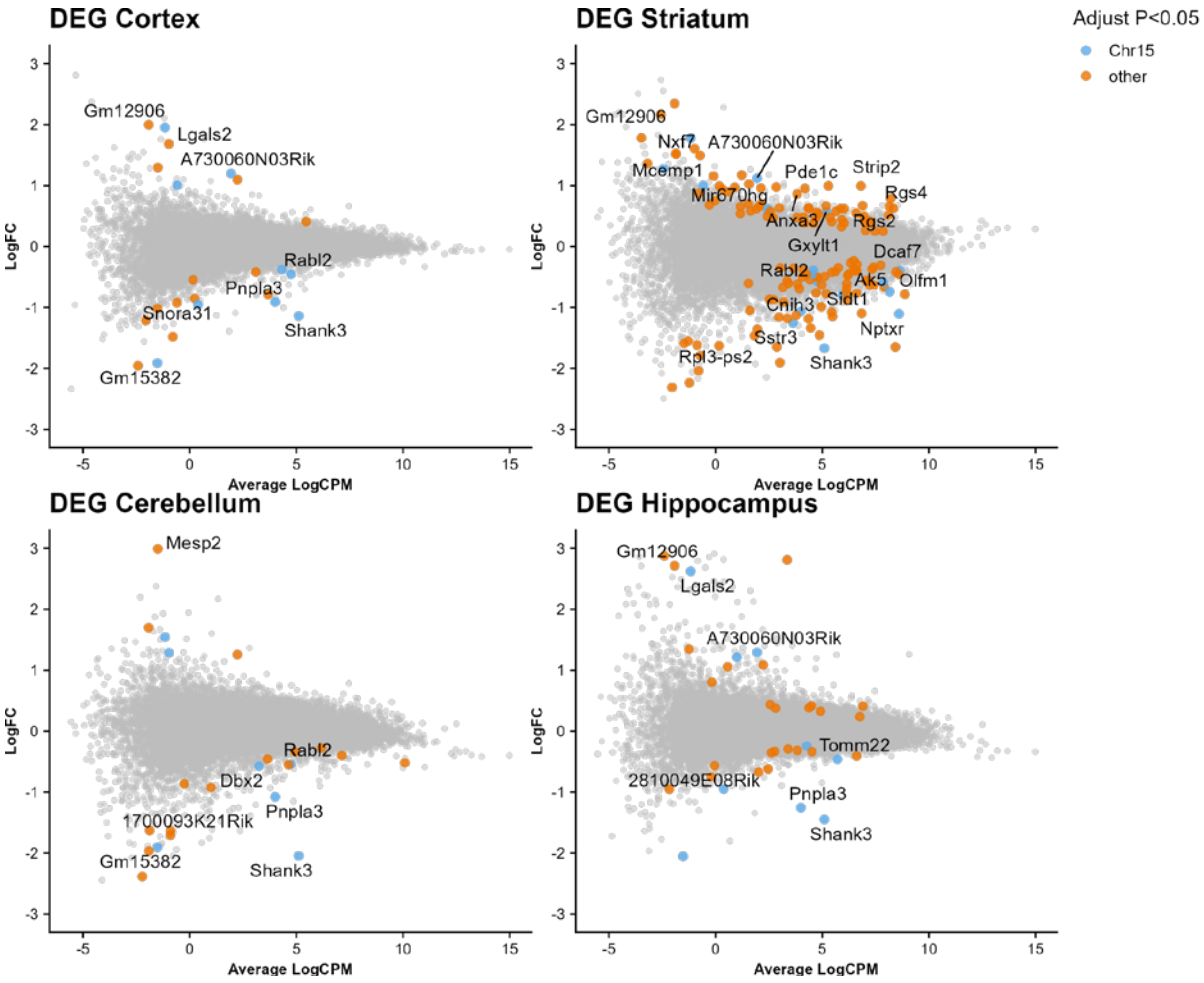


### Supplementary Figure S8

Gene expression differences in four brain regions of *Shank3*$^{\Delta11/\Delta11}$ mice compared to *Shank3*$^{+/+}$ mice. Mean-difference plots showing $\log_2$FC between *Shank3*$^{\Delta11/\Delta11}$ and *Shank3*$^{+/+}$ samples as a function of logCPM in the cortex (upper left), striatum (upper right), cerebellum (lower left) and hippocampus (lower right). Genes with a FDR lower than 5% are indicated by a coloured dot, blue for the ones located in the *Shank3* region on chromosome 15, and orange for the others. The star (*) indicates a pseudogene of a gene close to *Shank3* on chromosome 15. Only the names of the genes with an absolute log fold-change value larger than 1 are indicated.

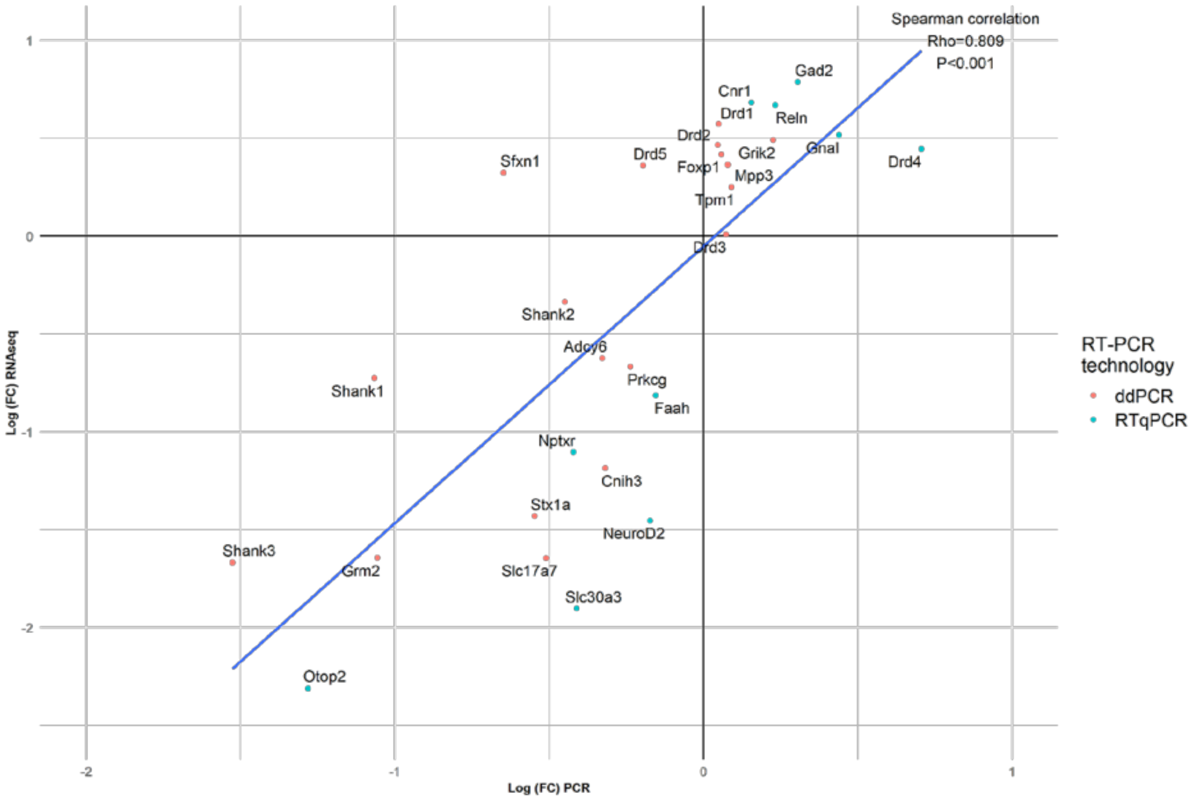


**Supplementary Figure S9**

Validation by quantitative RT-PCR of DEGs identified by RNAseq analysis in the striatum. Differential striatal expression between *Shank3*$^{+/+}$ and *Shank3*$^{\Delta 11/\Delta 11}$ mice of genes selected among the DEGs identified by RNA sequencing, as well as *Shank1*,*Shank2, and Drds* genes, using either the qRT-PCR (in blue) or dd-PCR (in red) technology.

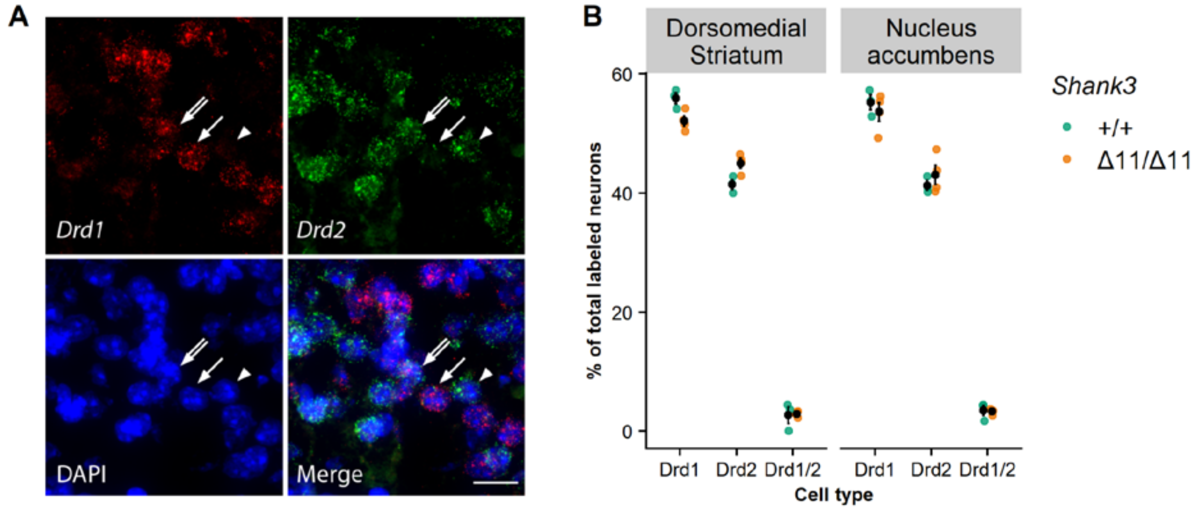


**Supplementary Figure S10**

The proportion of D1-MSN and D2-MSN is not affected in the *Shank3*$^{Δ11/Δ11}$ mice. (A) smFISH staining of D1 (*Drd1*, red) and D2 (*Drd2*, green) dopamine receptors transcripts. Nuclei are stained by DAPI (blue). Bright points represent RNA staining. White arrows point to the cell body of a D1-MSN, arrowheads to the cell body of a D2-MSN, and double arrows to the cell body of a D1/D2-MSN. (B). Proportion of nuclei associated with the *Drd1* or/and *Drd2* staining in the dorsomedial striatum and in the nucleus accumbens core in *Shank3*$^{+/+}$ mice (green) and *Shank3*$^{Δ11/Δ11}$ mice (orange). Green and orange points represent individuals (10-15 images per individual) and black points are means ± s.e.m.

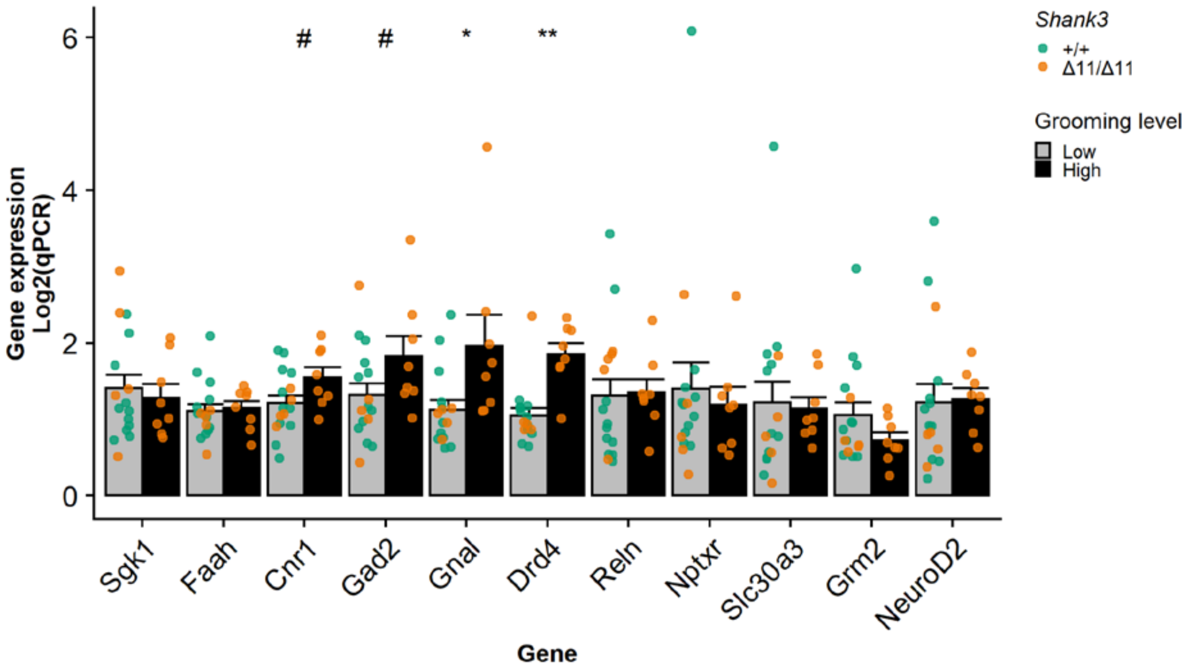


**Supplementary Figure S11**

Gene expression in mice with low or high self-grooming. Striatal expression of genes (RT-q-PCR) selected among the DEGs identified by RNA sequencing in mice with low or high self-grooming. Mann–Whitney U test: #: p<0.1; *: p<0.05; **: p<0.01.

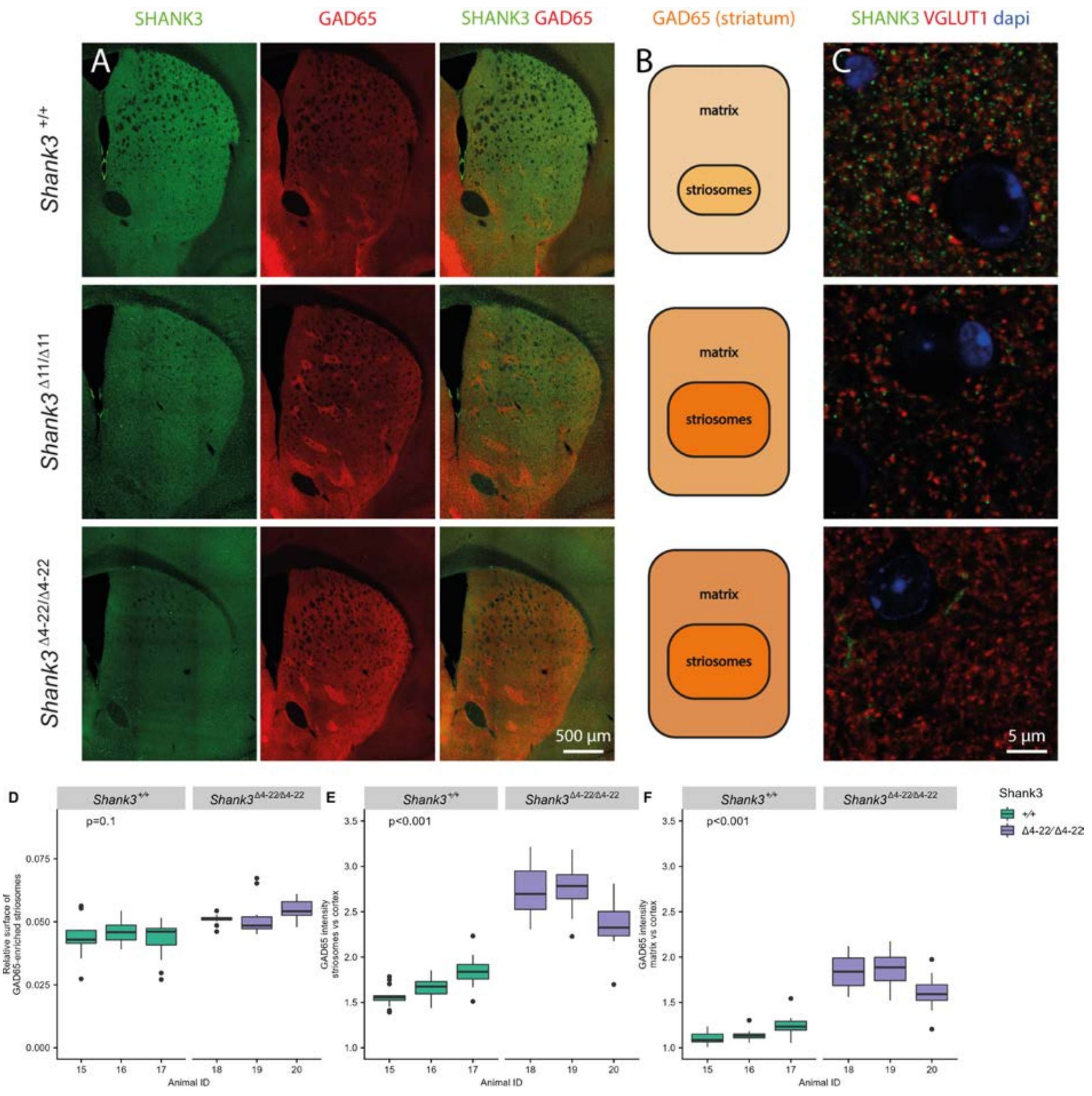


**Supplementary Figure S12**

Over-expression of GAD65 and enlargement of the striosomal compartment in the striatum of two different SHANK3-deficient mouse models, *Shank3*$^{\Delta11}$ and *Shank3*$^{\Delta4\text{-}22}$. (A) SHANK3 and GAD65 in the striatum. Images were generated by stitching multiple confocal images of coronal brain sections immunostained for SHANK3 and GAD65 (green in left panels and red in centre panels, respectively, merged images in right panels) in *Shank3*$^{+/+}$ (upper panels), *Shank3*$^{\Delta11/\Delta11}$ (middle panels) and *Shank3*$^{\Delta4\text{-}22/\Delta4\text{-}22}$ (lower panels) mice. Note that SHANK3 immunoreactivity is comparable in striosomes and matrix. (B) Schematic representation of GAD65 over-expression (colour gradient from light to dark orange) and of the enlargement of the striosomal compartment in the striatum of the *Shank3*$^{\Delta11/\Delta11}$ (middle panel) and the *Shank3*$^{\Delta4\text{-}22/\Delta4\text{-}22}$ (lower panel) mice, compared to *Shank3*$^{+/+}$ mice (upper panel). GAD65 overexpression is much greater in the striosomes than in the matrix in *Shank3*$^{\Delta11/\Delta11}$ mice that still express some SHANK3 isoforms, while in the *Shank3*$^{\Delta4\text{-}22/\Delta4\text{-}22}$ complete knock-out mice, GAD65 over-expression is great in both compartments. (C) Specificity of the anti-SHANK3 antibody. As expected for a protein of the post-synaptic density of the glutamatergic synapse, in the *Shank3*$^{+/+}$ mice (upper panel), SHANK3-positive puncta (green) are adjacent to the red puncta revealing VGLUT1, a marker of the pre-synaptic density of the glutamatergic synapse. A residual specific staining is still observed in the *Shank3*$^{\Delta11/\Delta11}$ mice (middle panel), while no synaptic puncta is observed in the *Shank3*$^{\Delta4\text{-}22/\Delta4\text{-}22}$ mice (lower panel). Nuclei are stained in blue by DAPI. (D, E, F) Comparison of GAD65 immunoreactivity in the striosome and matrix compartments of the dorsal striatum in 3 *Shank3*$^{+/+}$ (green) and 3 *Shank3*$^{\Delta4\text{-}22/\Delta4\text{-}22}$ (orange) 20-28 weeks old male mice. (D) Relative surface of the GAD65-enriched striosome compartment (surface of striosomes / surface of (striosomes + matrix). (E) Relative GAD65 labelling intensity in the striosomal compartment of the striatum compared to the cortex. (F) Relative GAD65 labelling intensity in the matrix compartment of the striatum compared to the cortex. Data, generated from analysis of at least 11 images per animal, are presented as box-plots (median, first, and third quartiles).

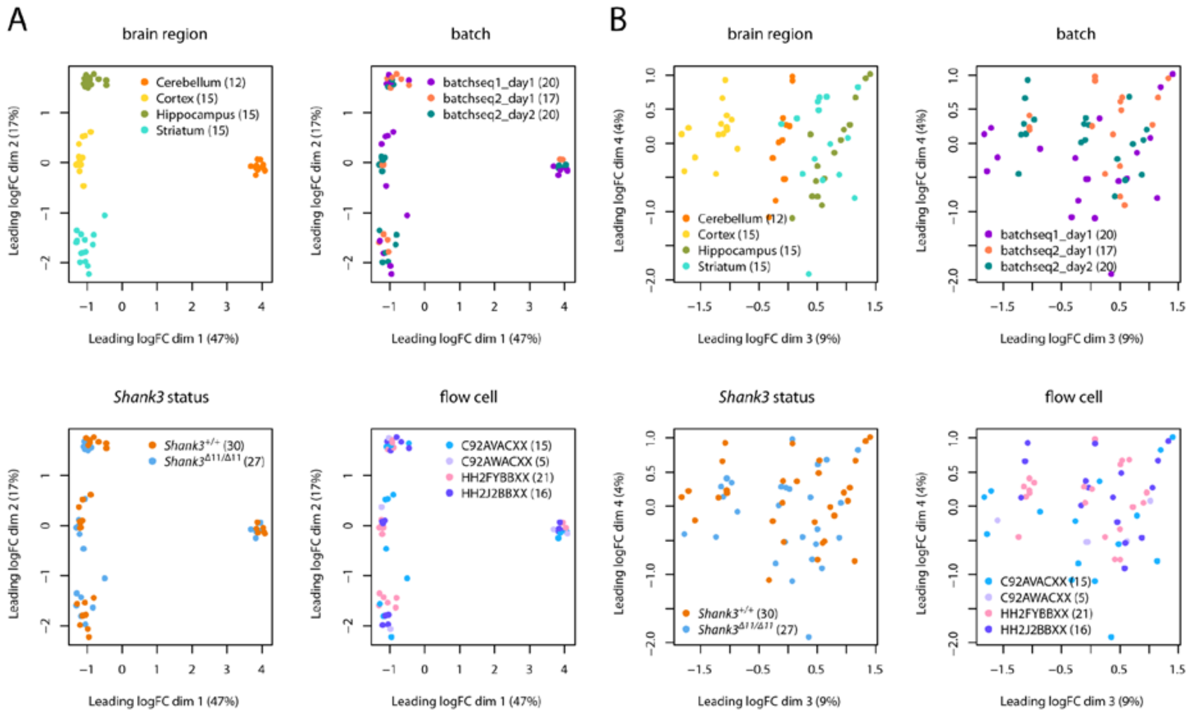


### Supplementary Figure S13

Multidimensional scaling (MDS) plots of the gene log counts-per-million (logCPM). (A) Data showing the positions of the samples in the space spanned by the first and second MDS dimensions. Samples are coloured with respect to brain region (upper left), batch of sequencing and RNA extraction (upper right), genotype (lower left), and flow cell (lower right). (B) Data showing the positions of the samples in the space spanned by the third and fourth MDS dimensions. Samples are coloured with respect to brain structure (upper left), batch of sequencing and RNA extraction (upper right), genotype (lower left), and flow cell (lower right).

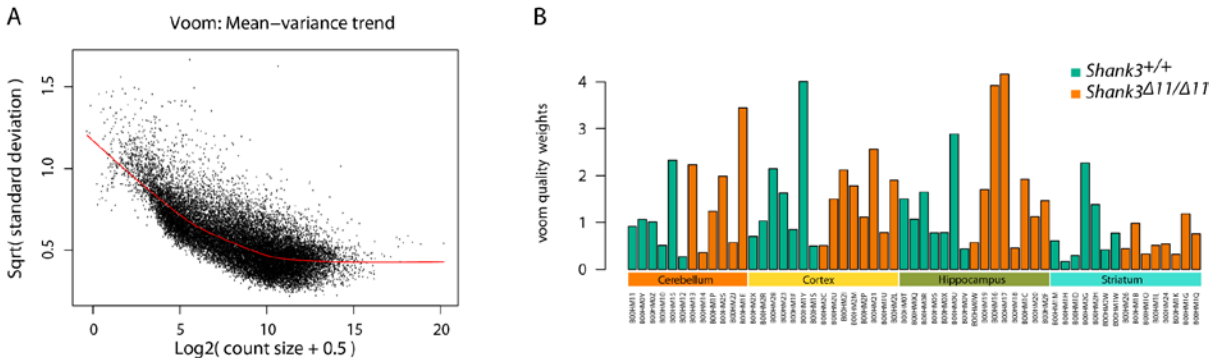


**Supplementary Figure S14**

Mean-variance relationship. (A) Voom mean-variance trend output for the RNAseq data. Gene-wise means and variances of the RNAseq data are represented by black points with a LOWESS trend. (B) 'Voom' sample-specific weights. Each bar plot represents the weight of one sample used to model variability between different samples.

### Supplementary material and methods

Animals and *Shank3*$^{\Delta 11}$ cohorts

Mice were weaned at 22 ± 1 days of age and housed in same-sex mixed-genotype groups of three to five animals unless otherwise specified. Mice were housed under constant temperature (22 ± 1 °C) with a 12:12 light/dark cycle (light on from 07:00 AM to 07:00 PM).

Cohort 1 involved 44 males (13 *Shank3*$^{+/+}$, 19 *Shank3*$^{+/\Delta 11}$ and 12 *Shank3*$^{\Delta 11/\Delta 11}$) and 38 females (10 *Shank3*$^{+/+}$, 16 *Shank3*$^{+/\Delta 11}$ and 12 *Shank3*$^{\Delta 11/\Delta 11}$). Mice from Cohort 1 were evaluated at Institut Pasteur at 3 months of age -the standard age for behavioural testing- for dark-light, Y-maze, open field tests, self-grooming observation, 3-chambered test, free social interactions, as well as at 8 and 12 months of age for open field test, self-grooming and free social interaction. After the tests at three months of age, males were housed individually, while females remained group-housed. Cohort 2 involved 42 males (15 *Shank3*$^{+/+}$, 14 *Shank3*$^{+/\Delta 11}$ and 13 *Shank3*$^{\Delta 11/\Delta 11}$) and 22 females (4 *Shank3*$^{+/+}$, 11 *Shank3*$^{+/\Delta 11}$ and 7 *Shank3*$^{\Delta 11/\Delta 11}$). Male and female mice from Cohort 2 were evaluated at Institut Pasteur at three months of age for self-grooming observation, open field, elevated plus maze, dark-light anxiety and rotarod tests. Cohort 3 involved 14 *Shank3*$^{+/+}$ and 7 *Shank3*$^{\Delta 11/\Delta 11}$ males that were bred at Institut Pasteur and sent at 2-2.5 months to Institut Biologie Paris Seine (IBPS). They were subjected to the open field, elevated plus maze, rotarod, Morris water maze and starmaze tests.

Behavioural tests

*General health*

At weaning, we measured weight and observed hind limb clasping as well as the physical aspect (fur, injury, or malformation) of the mice. At three and twelve months of age in Cohort 1, we again measured weight and noted the physical aspect of all mice.

*Dark/light anxiety-like test*

The mouse was left to freely explore a test cage separated into two compartments connected by a small door (5x5 cm; dark side: 3 lux; light side: 1300 lux) for 5 minutes. The latency to enter the dark compartment and the time spent in each compartment were measured (Ethovision, Noldus

Information Technologies, Wageningen, The Netherlands). More anxious mice are expected to spend shorter time in the light compartment and do less transitions between the compartments.

Y-maze

The animal (Cohort 1) was allowed to freely explore a y-maze during 5 min. The number of entrances as well as the sequence of entrances into each arm were manually noted by an experimented scientist. The sequence of good alternations, i.e. when the subject does not go back to the previous arm visited, are used to determine the working memory.

*Elevated plus-maze anxiety-like test*

The animal (Cohort 2) was allowed to freely explore the setup for 10 min. The elevated plus-maze (four arms of 7 cm by 30 cm, 50 cm above the floor) consisted of two open arms (no walls), and two closed arms (with walls), all connected by a neutral zone in the centre (100 lux). We measured the time spent in the open and closed arms, as well as the number of transitions using Ethovision (Noldus Information Technologies, Wageningen, The Netherlands). The more anxious the mice are the more time they spend in the closed arms. Cohort 3 was assessed at IBPS using a cross-shaped maze made of black perspex with a central zone (8 x 8 cm) facing closed and open arms (24 x 8 cm, surrounded by 25 cm walls made of grey perspex), elevated to a height of 50 cm. The percentage of time spent and the number of entries in the open arms was measured. An entry was considered valid when the 4 paws were present in the arm. The test lasted 5 min.

*Locomotion and exploratory test in the open field*

The mouse (Cohorts 1 & 2) was allowed to freely explore for 30 min a round open field arena of 1 m in diameter (100 lux in the centre of the arena). We recorded the total distance travelled (Ethovision, Noldus Information Technology, Wageningen, The Netherlands). Spontaneous locomotor activity of Cohort 3 was quantified in an arena made of grey perspex (45 x 45 cm) surrounded by red Plexiglas walls (30 cm height). Mice were first positioned in the centre of the arena and were allowed to freely explore for 10 min. Data acquisition was performed at a frequency of 25 Hz using the SMART® video recording system and tracking software and travelled distance was computed using NAT (Navigation Analysis Tool), a custom Matlab-based software [1].

*Observation of stereotyped behaviour*

Mice were individually placed in a new test cage (Plexiglas, 50 x 25 x 30 cm; 100 lux; clean sawdust bedding) in a soundproof chamber. After 10 min habituation, we recorded their behaviour for 10 min (camera Logitech C920). We manually scored the time spent self-grooming and digging (The Observer, Noldus Information Technology). When scoring self-grooming, we did not take into account the scratching behaviour. The total duration and the average event duration of each behavioural category were calculated.

*Occupant/new-comer social test with ultrasonic vocalisation recording*

The tested mouse was isolated socially for three days (females) or three weeks (males) to increase motivation for affiliative social contacts [2]. From this test on at 3 months of age, males remained singly housed. After 20 min of habituation for the tested mouse (the occupant) to the test cage (Plexiglas, 50 x 25 x 30 cm; 100 lux; clean sawdust bedding) in a soundproof chamber, an unfamiliar age- and sex-matched C57BL/6J mouse (new-comer, NC; Charles River Laboratories, France) was introduced. The two mice were allowed to freely interact for 4 min. Social interactions (video camera Logitech C920, 30 fps) were semi-automatically analysed using the 2D tracking module Mice Profiler from the ICY platform [3,4](Institut Pasteur, Paris, France). We quantified, for both the occupant and the new-comer, the time spent in contact, the types of contact (oral-oral contact, oro-genital contact), the "approach then escape" sequences, the follow behaviour, and the time spent in the vision field of the other one [5]. At the same time, ultrasonic vocalisations (USV) were recorded (Condenser ultrasound microphone Polaroid/CMPA, UltraSoundGate 416-200, Avisoft Bioacoustics, Glienicke, Germany; sampling frequency: 300 kHz; FFT-length: 1024 points; 16-bit format). Vocalisation files were analysed automatically using the vocalisation analysis plugin from LMT USV Toolbox [6].

*Male behaviour in presence of an oestrous female*

The tested male was placed in the presence of a female for 48 hours. Then, the male was isolated again for one day. During the test, the male was placed in the test cage (Plexiglas, 50 x 25 x 30 cm; 100 lux; with clean sawdust bedding) during 10 min for habituation [5]. After this period, an unknown C57BL/6J female in oestrus (tested through vaginal smears in the morning) was introduced into the test cage for 4 min. Both mice were allowed to freely interact. Social interactions as well as USV were recorded, as described above.

*Three-chambered social test*

A Plexiglas cage was divided in three connected compartments (side compartments: 150 lux; central compartment: 140 lux) as previously described [7]. Both side compartments contained an empty wire cup. First, the tested mouse was allowed to freely explore the setting, with doors open for 10 min (phase 1) for habituation. Then, the mouse was restricted in the central compartment, while an unfamiliar C57BL/6J mouse of the same sex (stranger 1) was placed under one of the cups (sides alternated between each mouse). The tested mouse was then allowed to explore the apparatus for 10 min (phase 2). In all phases, the time spent in each compartment and the number of transitions between compartments were automatically recorded. The time spent in contact with the cup containing the mouse (stranger 1) and the time spent in contact with the empty cup were manually measured in phase 2 to evaluate social interest.

*Morris Water Maze with Visible Platform*

Mice were trained in a circular water tank (150 cm in diameter, 40 cm high) to swim toward a visible platform marked with an object (11 cm high). The platform was randomly placed at different locations across trials and the pool was surrounded by blue curtains to occlude extramaze cues. Training consisted in one training session per day, four trials per session, during 2 days. The starting position (North, East, West, or South) was randomly selected with each quadrant sampled once a day. At the beginning of each trial, the mouse was released at the starting point and made facing the inner wall. Then, it was given a maximum of 90 s to locate and climb onto the escape platform. If the mouse was unable to find the platform within the 90 s period, it was guided to the platform by the experimenter. In either case, the mouse was allowed to remain on the platform for 30 s. Data acquisition was performed at a frequency of 25 Hz using the SMART® video recording system and tracking software.

*Starmaze*

The starmaze consisted of five alleys radiating from the vertices of a central pentagonal ring. All the alleys were filled with water, and the water was made opaque with an inert nontoxic product (AccuScan OP 301, Brenntag). The maze was surrounded by a square black curtain with 2D and 3D patterns affixed to provide configurations of spatial cues. To avoid the possible use of a guidance strategy (i.e., animals could rely on the use of a single distal cue), cue was given in duplicate. White noise was used to cover all other sounds that the mice could have used to orientate themselves. To solve the task, animals had to swim to a platform hidden 1 cm below the water surface and located 10 cm from the end of one alley. Departure and arrival points were always the same. All animals ran one session of five trials per day over 2 days using a 40-min inter-trial interval. If an animal did not locate the escape platform within 90 s, the experimenter placed the animal onto the platform for 30 s. During the protocol, one central alley and two peripheral alleys were blocked, forcing the mice to use the "left pathway." Mice were tracked by using the Smart Software (Bioseb, Vitrolles, France).

<u>Transcriptome analysis</u>

*Tissue collection*

Mice were killed during the light phase, specifically between 9:00 and 11:00 AM, by $CO_2$ intoxication. The brain was removed and macro-dissected on ice (4°C) into HBSS solution by an experienced practitioner. After separation of the hemispheres, six brain structures were extracted: whole cortex, hippocampus, whole striatum, cerebellum, diencephalon, and brainstem. Samples were snap frozen in liquid nitrogen and stored at -80°C.

*Total RNA extraction and sequencing*

Total RNA was extracted from four over six brain structures (whole cortex, hippocampus, whole striatum, cerebellum) using the miRNeasyPlus Micro Kit (Qiagen), following the manufacturer's instructions, including DNAse digestion. After first quality assessment using the Nanodrop spectrophotometer ND-1000 (Thermoscientific), the samples were analysed by the CNRGH (Centre National de Recherche en Génomique Humaine, CEA, Evry, France). RNA integrity was assessed using the Bioanalyzer RNA 6000 Nano assay and 2100 Bioanalyzer (Agilent Technologies). Then, an oriented mRNA sequencing was performed on samples with a RNA Integrity Number (RIN) larger than eight. Eight *Shank3*$^{\Delta 11/\Delta 11}$ and 7 *Shank3*$^{+/+}$ 12-month old mice were used for the differential expression analysis. For each of these mice, we studied four brain regions (cerebellum, cortex, hippocampus, and striatum), except for one in each group for which only three of the four brain regions were available. The description of the samples is available in Supplementary Table S5. The data were obtained from two batches of sequencing performed at eight months of interval. For the first batch, RNA extractions were performed on the same day while for the second sequencing batch, RNAs were extracted on two different days. We refer to this variable containing three levels as the batch variable. Each sequencing run included two flow cells.

*Mapping and reference genome*

The RNAseq reads were mapped onto the genome with the STAR aligner v2.5.3a [8] in 2-pass mode to a masked version of the *Mus Musculus* GRCm38 genome. During a first round of the differential gene expression analysis, we observed enrichment of the DEGs in genes located on the same arm of chromosome 15 as *Shank3*. Differences in genetic backgrounds around the *Shank3*$^{+/+}$ (C57BL/6J) and the *Shank3*$^{\Delta 11/\Delta 11}$ alleles (129S1/SvImJ, from the ES cells used to generate the *Shank3*$^{\Delta 11}$ mutation followed by 15 backcrosses on C57BL/6J) could affect the mapping by impacting the mapped reads on the remaining region of 129S1/SvImJ around *Shank3*. To avoid this bias, we masked the GRCm38 genome for variants of the 129S1/SvImJ mouse strain before mapping the sequencing reads. Variants were extracted from the VCF file provided by The Mouse Genome Project [9] (ftp://ftp-mouse.sanger.ac.uk/REL-1505-SNPs_Indels/mgp.v5.merged.snps_all.dbSNP142.vcf) and masked in the reference genome using the SNPsplit software (Krueger and Andrews 2016).

*Calling variants from the RNAseq data*

To see whether differences between the C57BL/6J and the 129S1/SvImj genomes in the region around *Shank3* could explain the larger number of DEGs detected in the vicinity of *Shank3*, we used the RNAseq data to identify single nucleotide polymorphisms (SNPs) in each mouse. We followed the Gatk Best Practices [10] workflow for SNP and indel calling on RNAseq data (https://software.broadinstitute.org/gatk/guide/article?id=3891) which includes the following steps: (i) map to the reference genome with STAR in multi-sample 2-pass mode to get the most sensitive novel junction discovery; (ii) add read groups, sorting, marking duplicates, and create index, using Picard's tools (http://broadinstitute.github.io/picard); (iii) split reads into exon segments (removing Ns but maintaining grouping information) and hard clipping sequences overhanging into the intronic regions, using the SplitNCigarReads Gatk tool; (iv) realign indels and recalibrate Base quality; (v) call variant with HaplotypeCaller, and finally filter the variants with VariantFiltration.

The last step was adapted to our project where several samples coming from the same mouse were available. HaplotypeCaller (with parameter -ERC BP_RESOLUTION) was called for each mouse individually using as input the processed BAM files coming from different brain tissues of the same mouse. Mice with the same *Shank3* status were then genotyped together by inputting the GVCF files to the Gatk tool GenotypeGVCFs. *Shank3*$^{\Delta 11/\Delta 11}$- and *Shank3*$^{+/+}$-specific VCF files were finally combined with the Gatk tool CombineVariants, which provides allele frequency and specificity of the variants to *Shank3*$^{\Delta 11/\Delta 11}$ or *Shank3*$^{+/+}$ mice populations. The resulting VCF file was filtered using the Gatk tool VariantFiltration using the parameters recommended in the Gatk workflow.

*Differential Expression Analysis*

The 18,194 genes with at least one count-per-million (CPM) in two samples were selected. The samples flagged by the QC Analyzer were filtered out. MDS plots showed separation of the samples according to brain regions, batches, and flow cells (**Supplementary Figure S13**).

Differential gene expression (DGE) analysis was performed with limma-voom v3.34.8 [11], and the version of *voom* using sample-quality weights[12] (function *voomWithQualityWeights*) in order to take into account the sample heterogeneity observed within and across brain regions (**Supplementary Figure S14**). The Trimmed Mean of *M*-values (TMM) method was used to calculate normalisation factors between samples. Three factors were included in the design matrix: the batch, the flow cell, and the brain tissue and *Shank3* status combined into one factor of eight levels. Since we were making comparisons both within and between mice, we treated the mouse as a random effect to adjust for baseline differences between subjects. To do so, the mouse was used as a blocking factor and the correlation between measurements made on the same mouse was computed using the function *duplicateCorrelation* and was input into the linear model fit, as suggested in the section "Multi-level Experiments" of the limma user guide. For each contrast of interest, the linear model was fitted for each gene using the function *lmFit*, and empirical Bayes smoothing was applied to the standard errors using the function *eBayes* with robust mode set to TRUE.

Gene-set over-representation analyses were performed using Fisher's hypergeometric tests. Plots were made using the R packages ggplot2[13], upsetR[14], and ggbio[15].

<u>Gene set and protein-protein interaction network analysis</u>

*Gene set analysis*

Gene set over-representation analyses were performed with the egsea.ora function available in the Bioconductor R package EGSEA v1.6.1 [16] without considering the genes around the *Shank3* gene. All collections of the databases MSigDB [17], GeneSetDB [18], and KEGG[19] were used.

*Protein-protein interaction network analysis*

Using the BioGRID database for *Mus Musculus* (https://thebiogrid.org/) [20], we analysed, for the 4 brain structures independently, the protein-protein interaction (PPI) network of the DEGs. The majority of DEGs for each structure was not annotated, i.e., not associated with a known pathway in BioGRID database (fraction of annotated DEGs: striatum: 71/186; hippocampus: 7/33, cerebellum: 4/24, cortex: 3/22). Cystoscape v3.6.1 was used to visualise the PPI network to find out key genes. A randomised protein interaction network was created using all DEGs to determine the probability of finding a network.

Quantitative RT-PCR

For dd-PCR, total RNA was extracted as described above and the cDNA library was generated using the iScript advanced cDNA kit (Bio-Rad). The dd-PCR was performed with the dd-PCR supermix for probes (no dUTP, Bio-Rad) and probes labelled with the FAM (for the genes of interest) and HEX (for the *Gapdh* housekeeping gene) fluorophores using the QX100 droplet digital PCR system (Bio-Rad). Results were analysed using the QuantaSoft Software.

For q-PCR, total RNA (200 ng) was reverse-transcribed with oligo-dT primers using RevertAid First Strand cDNA Synthesis Kit (ThermoFisher). Quantitative PCR was performed in triplicate with a 7500 real time PCR system (Applied Biosystems) using LightCycler SYBR Green I Master Mix (Roche) and specific pairs of primers. Individual data were normalised using a combination of two housekeeping genes (*Ppia* and *Rpl13a*). The results are reported as fold changes.

Single-molecule Fluorescent In Situ Hybridisation and immunofluorescence experiments on brain sections

*Single-molecule Fluorescent In Situ Hybridisation (smFISH) and image analysis*

Three month-old mice, deeply anesthetised by an intraperitoneal injection of a mixture of Ketamine (Imalgen®, 200 mg/kg, Merial) and Xylazine (Rompun®, 8 mg/kg, Bayer), were transcardially perfused with 10 ml of PBS, followed by 50 ml 4% paraformaldehyde (PFA) in PBS (Santa Cruz Biotechnology) at 4°C. Each brain was snap frozen in liquid nitrogen and then stored at -80 °C. Cryosections (16 µm thick), obtained using the CM300 cryostat (Leica Biosystem) were collected on Superfrost Plus microscope slides (Thermo Fisher Scientific) and stored at −20°C before hybridisation. The brain section (from bregma 1.7 to -0.58) were treated as described by Tsanov et al. (2016) [21] using 48 probes along *Drd1* RNA and 40 probes along *Drd2* RNA (see **Table S6**). Images were acquired in the dorso-medial striatum and in the nucleus accumbens core regions using an Axio observer Z1 inverted microscope (Carl Zeiss) equipped with a Plan-Apochromat 20X/0,8 M27 objective (Carl Zeiss). Images were manually analysed using imageJ software (NIH).

*Confocal fluorescence microscopy on brain sections*

Immunofluorescence analyses on striatum sections were performed on one year old (52-70 weeks) *Shank3*$^{\Delta 11}$ and 20-28 weeks old *Shank3*$^{\Delta 4\text{-}22}$ male mice. Animals, deeply anesthetised as described above, were transcardially perfused with 20 ml of PBS, followed by 50 ml 4% paraformaldehyde (PFA) in PBS (Santa Cruz Biotechnology). Brains were removed and fixed overnight at 4°C in 4% PFA, rapidly washed in PBS, and then immersed in 15% sucrose in PBS for overnight incubation at 4°C. The 15% sucrose solution was then replaced by a 30 % sucrose solution for a second overnight incubation at 4°C. Brains were then transferred in Shandon Cryomatrix™ Frozen Embedding Medium (Thermo Scientific™) in cryomolds and frozen by immersion in 2-methyl butane (Sigma-Aldrich) chilled in liquid nitrogen. Samples were kept at -80°C until use. Forty µm coronal sections, obtained using the CM3050S cryostat (Leica Biosystems), were rapidly transferred into PBS in 12-well-culture plates. The samples for GAD65 immunostaining were collected every five sections and grouped together in a separate single well and stored in PBS at 4°C until use. The free-floating sections were rinsed 3 times in PBS (pH 7.4) and then incubated in NH4Cl (Sigma-Aldrich) 50mM in PBS for 15 min. After three PBS washes (5 min each), sections were incubated for one hour at room temperature in PBS containing 1% bovine serum albumin (Applichem) and 0.3% Triton X-100 (Sigma-Aldrich) (PBS/BSA/TX) before incubation for ≈ 20 hours at room temperature with the primary antibodies (rabbit polyclonal anti-GAD65 (Invitrogen PA5-77983, 1/200), mouse monoclonal anti-GAD65 (Millipore MAB351, 1/500), guinea pig polyclonal anti MOR (Millipore AB5509, 1/100), mouse monoclonal anti-SHANK3 (Santa Cruz Biotechnology sc-377088, 1/1000), rabbit polyclonal anti-VGLUT1 (Synaptic Systems 135303, 1/1000) in PBS/BSA/TX. After three PBS washes (10 min each), sections were incubated in secondary antibodies (Alexa fluor 555™-conjugated goat anti-rabbit IgG, Alexa fluor488™-conjugated goat anti-mouse IgG and Alexa fluor488™-conjugated goat anti-guinea-pig IgG (all from Invitrogen)) diluted 1/500 in PBS/BSA/TX for one hour at room temperature, washed in PBS (3 washes of 10 min) and mounted in FluorSave™ Reagent (Calbiochem). To stain nuclei, a 10 min incubation step in DAPI (1µg/ml in $H_20$, Thermoscientific) followed by a PBS wash was added before mounting. Images were acquired using a LSM 700 confocal microscope (Carl Zeiss) equipped with a Plan-Apochromat 10X/0,45 M27 objective (Carl Zeiss). For each animal, 5 to 7 coronal sections spanning the anterior-posterior axis from Bregma 1.10 mm to Bregma 0.14 mm were imaged bilaterally. Acquisition parameters were adjusted for each brain hemi-section in order to have no saturating signal for GAD65 in the striatal region of interest. Images of the dorsal striatum were reconstructed by stitching multiple maximum-intensity projected z-stacks.

*Quantitative analysis of GAD65 immunoreactivity*

To deal with sample to sample variations in labelling efficiency inherent to immunofluorescent labelling methods we determined, for each image, the ratio of the labelling intensities of the striosome and matrix compartments to the adjacent cortex, in which *Gad2* is not differentially expressed in Shank*3*$^{\Delta11/\Delta11}$ and *Shank3*$^{+/+}$ mice, according to the transcriptome analysis. For each image, the regions of interest (dorsal striatum and adjacent cortex) were delimited using the Icy software (Institut Pasteur, Paris). In case of edge effect along the lateral ventricle, the concerned edge was excluded from the region of interest. We developed a method to automatically detect three areas in the striatum: the unlabelled myelinated fibres, the matrix (lower expression of GAD65), and the patches/striosomes (higher expression of GAD65). To deal with possible acquisition artefacts, we first applied a mean filter on the image by using a box blur (size 7x7). We then pre-computed two maps corresponding to the local mean and standard deviation of the image. For each point, we computed those values considering a window of 1000x1000 pixels centred on the point, with a computation step of 10px. Then, for each pixel of the smoothed image, we labelled as fibres all points of intensity value below localMean-localStd. The final fibre mask was eroded morphologically (2px) to remove spurious segmentations and connectivities. To compute the patches, we performed the same computation, but the pixel intensity had to be over localMean+localStd and below localMean + localStd*5 to remove super-bright spots corresponding to fluorescent dust. We also eroded the result by a factor of 2 pixels. Fibber and Patch masks were then processed with a 2D 8-way connected component process to get all individual fibres and patches. We then cleaned up the detection by removing patches of a surface inferior to 100 pixels, and fibres of a surface inferior to 50 pixels. To determine the intensity ratio of matrix versus cortex, we created a matrix segmentation corresponding to the striatal region of interest subtracted by the patches and the fibres detected. The meanMatrix was computed over this surface while the meanPatch was the mean intensity of all pixels contained in the patch mask. For each image, we computed the ratio meanPatch over cortex and meanMatrix over cortex. We also computed the surface of the patch mask relative to the surface of the (patch mask + matrix mask). To determine the effect of genotype on the data extracted by the image analysis, we applied a linear mixed model (LMM) using the method "restricted maximum likelihood (REML)" from the Python library "statsmodels.formula.api.mixedlm". In all presented results, the LMM has converged and a p-value indicating if the datasets from the *Shank3*-mutated and *Shank3*$^{+/+}$ mice are significantly different is provided
